# MXRA8-dependent chikungunya virus infection in PDGFR ^+^ mesenchymal cells increases vascular permeability and facilitates hematogenous spread

**DOI:** 10.64898/2026.10.06.757040

**Authors:** Autumn C. Holmes, Yael Alippe, Deborah J. Lenschow, Michael S. Diamond

## Abstract

Alphaviruses are mosquito-transmitted pathogens in the *Togaviridae* family that cause severe arthritogenic or encephalitic disease in humans. Swelling of peripheral joints and soft tissues is a hallmark of arthritogenic alphavirus infection, although the mechanism by which tissue edema occurs is not well understood. The cell surface protein, matrix remodeling-associated protein 8 (MXRA8), is a cell entry receptor for several arthritogenic alphaviruses. Global loss of MXRA8 expression in mice abrogates foot swelling and reduces virus dissemination. Here, we demonstrate that specific ablation of MXRA8 from platelet-derived growth factor receptor alpha (PDGFRα) cells, including fibroblasts, phenocopies the foot swelling and dissemination deficits observed in chikungunya virus (CHIKV)-infected global MXRA8 knock-out mice. Edema in the foot requires CHIKV infection of PDGFRα^+^ cells, which enhances both infection efficiency and permeability of endothelial cells. CHIKV-induced vascular leakage requires both MXRA8 expression and vascular endothelial growth factor signaling and contributes to dissemination by enabling virus access to the bloodstream. Collectively, our study establishes the role of MXRA8-dependent cell tropism in CHIKV dissemination and pathogenesis.

## INTRODUCTION

Arthritogenic alphaviruses are emerging, mosquito-borne pathogens in the *Togaviridae* family that are maintained in urban or sylvatic transmission cycles and responsible for explosive outbreaks of debilitating arthralgia, myalgia, and musculoskeletal tissue swelling in humans. Although rarely fatal, joint and muscle symptoms can last for months to years in a subset of patients, resulting in prolonged disability. The pathology caused by arthritogenic alphaviruses, including chikungunya virus (CHIKV), is thought to occur because of their tropism for different cells in musculoskeletal tissues [1] particularly, those of mesenchymal origin. Muscle damage is attributed in part to infection of myofibers and myoblasts [2], and impaired bone mineralization and homeostasis may be due to infection of osteoblasts and chondrocytes [3–5]. CHIKV infection of fibroblast-like cells and synoviocytes elicits the recruitment of myeloid cells through the production of CCL2 [6], and these cells, along with CD4^+^ T cells [7, 8] sustain chronic synovitis and joint swelling [9].

A key determinant of virus cellular tropism is the expression of cognate entry receptors [10–12]. Several cell surface molecules have been described as attachment and entry factors for alphaviruses [10]. The cell surface protein, matrix remodeling-associated protein 8 (MXRA8), is a key entry receptor [13] for members of the Semliki Forest (SF) complex of alphaviruses, including CHIKV, Ross River virus (RRV), Mayaro virus (MAYV), and O’nyong’nyong virus (ONNV). MXRA8 is broadly expressed in fibroblasts, endothelial cells, mesenchymal cells, synoviocytes, and chondrocytes [14], Among its endogenous functions, MXRA8 acts as a cell adhesion molecule to facilitate leukocyte migration across the blood-brain barrier [15] and also promotes chondrocyte proliferation during embryogenesis [16]. Cell culture studies suggest that MXRA8 can inhibit angiogenesis and osteoclastogenesis by downregulating the p38 mitogen-activated protein (MAP) kinase pathway [16, 17]. In CHIKV, RRV, and MAYV-infected mice, global deletion of MXRA8 expression abrogates foot swelling near the site of inoculation and reduces virus dissemination to distant sites [14]. However, viral load at the inoculation site was largely unchanged in *Mxra8*^Δ*8/*Δ*8*^ global knockout (KO) mice [14]. This observation is consistent with data suggesting that CHIKV replication within skeletal muscle and the ensuing inflammatory response are uncoupled events [18].

Here, we generated conditional deletions of *Mxra8* in mice to better define its role in cell tropism, inflammation, and CHIKV pathogenesis. Deletion of MXRA8 expression in hematopoietic cells (*Vav1*-*Cre Mxra8^fl/fl^* mice) had no effect on CHIKV infection, foot swelling, dissemination after subcutaneous inoculation on the footpad. In contrast, deletion of MXRA8 in platelet-derived growth factor alpha (PDGFRα)-expressing mesenchymal cells (*Pdgfra*-*Cre^ERT2^ Mxra8^fl/fl^* mice) resulted in decreased footpad swelling at the site of CHIKV inoculation and reduced dissemination to distal musculoskeletal tissues. The phenotypes in *Pdgfra*-*Cre^ERT2^ Mxra8^fl/fl^* mice were associated with specific reductions in infection of fibroblast and adipocyte progenitors (FAPs) and endothelial cell populations, although overall changes in CHIKV viral burden in the ipsilateral foot were not detected compared to Cre- *Mxra8^fl/fl^* mice. Subsequent cell culture and *in vivo* experiments showed that direct infection of endothelial cells was MXRA8- independent, although efficient infection required MXRA8-dependent infection of fibroblasts. Finally, blockade of vascular endothelial growth factor (VEGF-A) reduced CHIKV-induced vascular permeability, foot swelling, and dissemination, and secretion of VEGF-A by FAPs in the context of CHIKV infection was MXRA8-dependent. Our data support a model in which MXRA8- dependent and -independent infection of FAPs and endothelial cells, respectively, act in concert to promote tissue edema, which provides the virus with access to the bloodstream, enabling greater viremia, more rapid dissemination to distant sites, and likely, enhanced transmission back to vector hosts.

## RESULTS

### Global MXRA8 deletion reduces CHIKV infection of CD45^-^ and CD45^+^ cell subsets without impacting overall viral burden in the foot

Although swelling of the foot proximal to the site of injection is reduced in global *Mxra8*^Δ*8/*Δ*8*^ KO mice, levels of virus are diminished by only 2-fold [14], suggesting that tissue edema and virus infection are not directly linked. To explore this hypothesis further, we assessed whether increasing the levels of infectious virus in the ipsilateral feet of *Mxra8*^Δ*8/*Δ*8*^ KO mice could rescue the swelling phenotype. Whereas raising the inoculating dose of CHIKV-AF15561 (CHIKV-AF) from 10^3^ to 10^5^ focus-forming units (FFU) restored virus titers to wild-type (WT) levels in the ipsilateral feet of *Mxra8*^Δ*8/*Δ*8*^ mice at 3 days post-infection (dpi, **Fig 1A**), foot swelling (**Fig 1B**) and viremia (**Fig 1C**) were not rescued.

**Figure 1.**
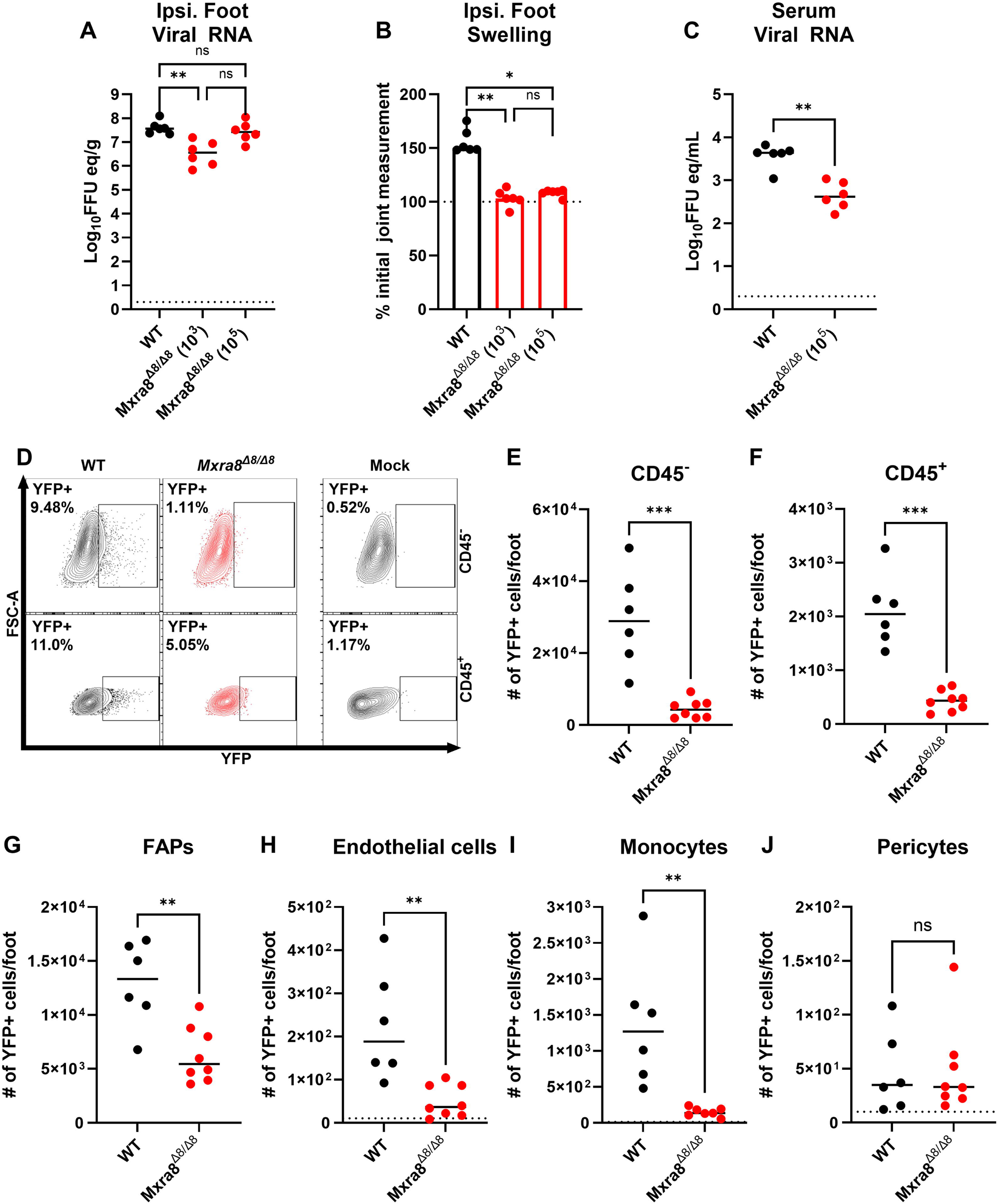
MXRA8-dependent infection of cell subsets in mice. (**A-B**) Cohorts of 4- week-old C57BL6/J WT mice (n = 6 mice per group, 2 experiments) were inoculated with 10^3^ or 10^5^ FFU of CHIKV-AF15561 in the left rear footpad (ipsilateral foot). (**A**) At 3 dpi, viral RNA was extracted from tissues and measured by qRT-PCR. Viral RNA is represented as FFU equivalents and is normalized to tissue weight. (**B**) At 3 dpi, footpad swelling was measured using digital calipers and is presented as the percentage of the initial weight measurement at day 0 (dotted line). (**C**) WT mice (n = 6 per group, 2 experiments) were inoculated subcutaneously with 10^3^ or 10^5^ FFU of CHIKV-AF15561 in the foot. At 3 dpi, serum was harvested, and viral RNA was measured. (**D-F**). Four-week-old C57BL6/J WT or *Mxra8*^Δ*8/*Δ*8*^ KO mice (n = 6-8 mice per group, 2 experiments) were inoculated with 10^5^ FFU of CHIKV-AF15561- Venus. At 2 dpi, the ipsilateral foot was harvested and digested to release stromal and hematopoietic cells. The percentage (**D**) and number of CD45^-^ (**E**) or CD45^+^ (**F**) cells expressing the Venus reporter protein (YFP^+^) were identified by flow cytometry. Gating strategies are in **Fig S1**. (**G-J**) Cell suspensions were also incubated with anti-CD31, anti-CD29, anti-Sca-1, anti- CD146, anti-CD11b, and anti-Ly6C antibodies to determine the number of YFP^+^ FAPs (**G**), endothelial cells (**H**), monocytes (**I**), and pericytes (**J**). Statistical analysis: (**A-B)** Kruskal-Wallis ANOVA with Dunn’s post-test; (**C-J**) Mann-Whitney test; *P\** < 0.05, \*\**P* < 0.01, \*\*\**P* < 0.001, ns, not significant. Dotted horizontal line on graph in **C** indicates limit of detection for the assay.

Skeletal muscle is comprised of myofiber bundles flanked by interstitial cell populations, including FAPs, endothelial cells, pericytes, and tissue resident macrophages [19], all of which are possible targets of infection by CHIKV and other arthritogenic alphaviruses [1, 20]. CHIKV RNA levels in muscle fibers of the ipsilateral feet are largely unchanged in the absence of MXRA8 [14]. However, labeling of infected fibroblasts in the musculoskeletal tissues of Cre- recombinase-inducible tdTomato reporter mice using a recombinant CHIKV-Cre recombinase virus was reduced by administration of anti-MXRA8 monoclonal antibodies (mAbs) [21], suggesting that fibroblasts, but not myocytes, require MXRA8 for efficient entry and infection. To better define MXRA8 tropism, we inoculated WT or *Mxra8*^Δ*8/*Δ*8*^ mice with a recombinant CHIKV strain that expresses a fluorescent protein (CHIKV AF-Venus [22, 23]). To ensure that endpoint viral titers in the ipsilateral feet were equivalent between WT and *Mxra8*^Δ*8/*Δ*8*^ KO mice, we inoculated the *Mxra8*^Δ*8/*Δ*8*^ KO mice with a higher dose of CHIKV AF-Venus (10^5^ FFU) and then harvested the ipsilateral foot at 2 dpi for flow cytometric analysis, a time point immediately prior to the first peak of foot swelling. We detected reduced (3-fold, *P* < 0.05) numbers of YFP^+^ cells in both CD45^-^ and CD45^+^ compartments in *Mxra8*^Δ*8/*Δ*8*^ KO compared to WT mice (**Fig 1D-F, S1**). Notably, FAPs (CD45^-^, CD31^-^, CD29^+^, Sca-1^+^), endothelial cells (CD45^-^, CD31^+^), and infiltrating monocytes (CD45^+^, CD11b^+^, Ly6C^hi^) showed a dependence on MXRA8 for CHIKV AF-Venus infection, whereas pericytes did not (**Fig 1G-J**). These data suggest that MXRA8 is required for infection of some cell types in the foot yet has a dominant role in contributing to infection-induced tissue swelling.

### CHIKV infection of PDGFR ^+^ stromal cells is MXRA8-dependent

To assess further the cell type-specific roles of MXRA8 in CHIKV infection and foot swelling, we generated cell-type specific deletions in newly generated *Mxra8*^fl/fl^ mice. We first crossed *Mxra8*^fl/fl^ mice to tamoxifen-inducible *Pdgfra*-*Cre^ERT2^* mice to conditionally delete MXRA8 from FAPs and their fibroblast progeny [24–27] in musculoskeletal tissues. To induce recombination in *Pdgfra*-*Cre^ERT2^ Mxra8^fl/fl^*mice, we administered 2 mg/ of tamoxifen per mouse every other day for 5 days (**Fig 2A**). We confirmed the efficiency of Cre recombination and *Mxra8* deletion by polymerase chain reaction (PCR) using primers that target the first 22 nucleotides of the *Mxra8* exon 1 and the last 22 nucleotides of the 3’ *loxP* site flanking *Mxra8* exon 9 (**Fig S2A**). We amplified an 800- basepair band in *Pdgfra*-*Cre^ERT2^ Mxra8^fl/fl^* mice but not Cre- mice, which corresponds to the residual *Mxra8* gene after excision of exons 2-9 (**Fig S2B**).

**Figure 2.**
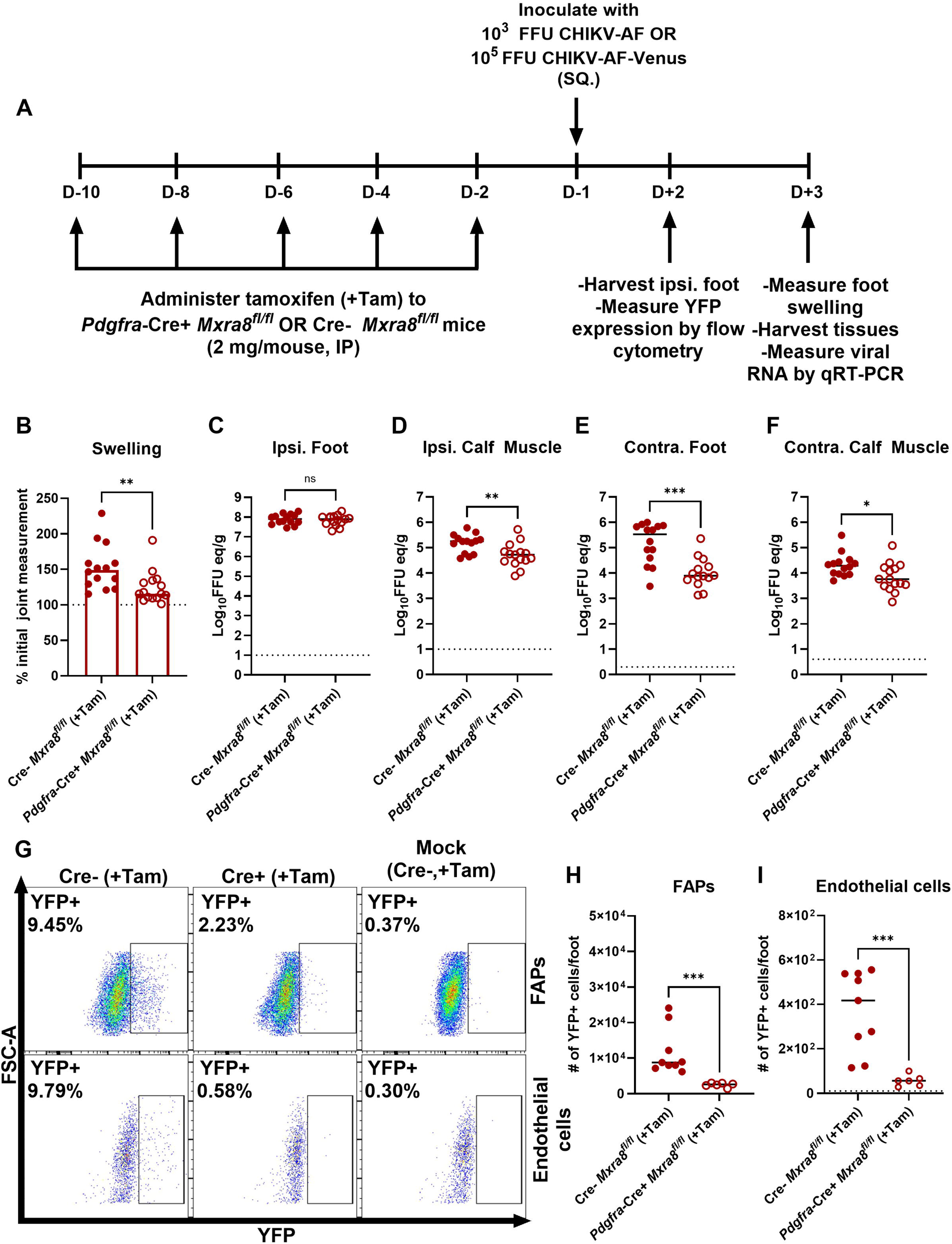
CHIKV infection in *Mxra8*^fl/fl^ conditional KO mice. (**A-G**) Three-week-old male and female *Pdgfra-Cre^ERT2^ Mxra8^fl/fl^*mice were administered tamoxifen (Tam) every other day for 5 days by intraperitoneal injection. *Pdgfra-Cre^ERT2^* or Cre- mice (n = 14-15 mice per group, 3 experiments) were inoculated with 10^3^ FFU of CHIKV-AF15561 via footpad injection. (**B**) At 3 dpi, footpad swelling was measured. Mice were then sacrificed, perfused with PBS, and ipsilateral foot (**C**), ipsilateral calf muscle (**D**), contralateral foot (**E**), and contralateral calf muscle (**F**) were harvested, and viral RNA was measured. (**G-I**) *Pdgfra-Cre^ERT2^* or Cre- mice were inoculated with 10^5^ FFU of CHIKV-AF15516-Venus in the footpad. At 2 dpi, ipsilateral feet were collected and processed as in Fig 1. The percentage (**G**) and number of FAPs (**H**) or endothelial cells (**I**) expressing YFP were quantified by flow cytometry. Statistical analysis: (**A-I**) Mann- Whitney test, *P\** < 0.05, \*\**P* < 0.01, \*\*\**P* < 0.001, ns, not significant. Dotted horizontal lines on graphs indicate limit of detection for the assay.

We inoculated tamoxifen-treated *Pdgfra*-*Cre^ERT2^ Mxra8^fl/fl^* mice, alongside tamoxifen-treated, Cre-negative (Cre-), *Mxra8^fl/fl^* littermate controls with 10^3^ FFU of CHIKV-AF. Deletion of MXRA8 in *Pdgfra*-*Cre^ERT2^ Mxra8^fl/fl^*mice resulted in diminished ipsilateral foot swelling (2-fold, *P* < 0.001) and reduced levels of viral RNA in the ipsilateral calf muscle (5-fold, *P* = 0.001), contralateral foot (50-fold, *P* = 0.0004), and contralateral calf muscle (5-fold, *P* = 0.012), but not in the ipsilateral foot (*P* > 0.05) (**Fig 2B-F**). Given that infection of FAPs was diminished in *Mxra8*^Δ*8/*Δ*8*^ KO mice, and swelling was reduced in *Pdgfra*-*Cre^ERT2^ Mxra8^fl/fl^* mice, we hypothesized that infection of FAPs was dependent on MXRA8 expression. To evaluate this idea, we inoculated tamoxifen-treated *Pdgfra*-*Cre^ERT2^* or Cre- *Mxra8^fl/fl^* mice with 10^5^ FFU of CHIKV-AF Venus and analyzed cells in the ipsilateral foot at 2 dpi by flow cytometry. YFP expression in FAPs and endothelial cells was reduced in Cre+ mice compared to the Cre- controls (**Fig 2G-I**). We also tested a second related arthritogenic alphavirus, Mayaro virus (MAYV) in our *Pdgfra*-*Cre^ERT2^ Mxra8^fl/fl^* model (**Fig S3A***-***S3F**). Like CHIKV, at 3 days-post-MAYV infection, ipsilateral foot swelling was reduced in *Pdgfra*-*Cre^ERT2^ Mxra8^fl/fl^* animals compared to Cre- controls (**Fig S3B**, 2-fold, *P* < 0.001). Viral RNA levels in the ipsilateral foot and ipsilateral calf muscle were not significantly different between the two groups (**Fig S3C-D**), but dissemination to contralateral sites was reduced in *Pdgfra*-*Cre^ERT2^ Mxra8^fl/fl^* mice compared to Cre- *Mxra8^fl/fl^* mice (**Fig S3E-F**: 5 to 10-fold, *P* < 0.001).

We also crossed *Mxra8*^fl/fl^ mice to *Vav1*-*Cre* mice to delete MXRA8 constitutively from hematopoietic [28–30] and endothelial cells [30, 31]. We validated the specificity and efficiency of our *Vav1*-*Cre Mxra8^fl/fl^* model and observed a band corresponding to the *Mxra8* excision product in blood from *Vav1*-*Cre Mxra8^fl/fl^* mice, which was largely absent in the tail (**Fig S4A**). Deletion of MXRA8 from hematopoietic cells or endothelial cell subsets in *Vav1*-*Cre Mxra8^fl/fl^* mice did not impact foot swelling, CHIKV RNA levels in the tissues we analyzed (**Fig S4B-F**), or YFP expression in FAPs or endothelial cells (**Fig S4G-I**). Together, these data suggest that arthritogenic alphaviruses may transit into the bloodstream from underlying infected musculoskeletal tissues, and that this dissemination mode requires MXRA8 expression in PDGFRα^+^cells.

In mammalian skeletal muscle, PDGFRα is predominantly expressed in FAPs [25, 32] and adopts a perivascular localization [33, 34]. However, a subset of skeletal muscle endothelial cells expresses *Pdgfra* mRNA [25], and PDGFRα^+^ CD31^+^ cells have been described [24, 35]. To determine if MXRA8 is deleted from endothelial cells in *Pdgfra*-*Cre^ERT2^ Mxra8^fl/fl^* mice, we sorted endothelial cells (CD45^-^, CD31^+^) and fibroblasts (CD45^-^, CD31^-^, CD29^+^) from the feet of uninfected, tamoxifen-treated *Pdgfra*-*Cre^ERT2^*or Cre- *Mxra8^fl/fl^* mice (**Fig S5A**) and measured *Mxra8* mRNA expression by qRT-PCR (**Fig S5B**). Deletion in *Pdgfra*-*Cre^ERT2^ Mxra8^fl/fl^* mice reduced (*P* < 0.05) *Mxra8* mRNA expression in fibroblasts, but not endothelial cells, relative to Cre- controls (**Fig S5B**). In a corroborating experiment, we tested if gene recombination occurs in fibroblasts or endothelial cells isolated from *Pdgfra*-*Cre^ERT2^ Mxra8^fl/fl^* mice after tamoxifen treatment (**Fig S5C**). From either Cre- or Cre+ fibroblasts or endothelial cells, we amplified the *Pdgfra*-*Cre* (lanes 1-4), the *Mxra8* gene excision product (lanes 5-8), and the 3’ *loxP* site (lanes 9-12) and (**Fig S5C**). The excision product band was absent in Cre- or Cre+ endothelial cells (lanes 11 and 12, **Fig S5C**). We also analyzed a published [36] single-cell RNA sequencing dataset to determine the cell types expressing *Mxra8* and *Pdgfra* in the mouse foot. *Pdgfra* transcripts were detected primarily in fibroblasts (72.5%), with < 6% in all other annotated populations. *Mxra8* transcripts were detected in fibroblasts (72.1%), stromal cells (36.4%) and endothelial cells (22.6%), with lower prevalence in macrophages (8.7%), and lymphoid and granulocyte populations (< 3.8%) (**Fig S5D**). These frequencies were comparable in mock and CHIKV-infected samples, indicating that infection does not alter the expression pattern of *Pdgfra* and *Mxra8* (**Fig S5E**). Together, these data establish that MXRA8 and PDGFRα are expressed principally in fibroblasts in the mouse foot, and that expression of *Mxra8* is reduced in fibroblasts but not in endothelial cells in *Pdgfra*-*Cre^ERT2^ Mxra8^fl/fl^* mice.

### MXRA8-dependent CHIKV infection of fibroblasts promotes endothelial cell infection and permeability

Because conditional deletion of MXRA8 in *Vav1-Cre Mxra8^fl/fl^* mice did not impact CHIKV infection (**Fig S4**), we hypothesized that endothelial cells might become infected by a MXRA8-independent pathway. To begin to address this question, we used two murine cell lines, NIH 3T3 fibroblasts and SVEC4-10 microvascular endothelial cells, that express similar levels of MXRA8 on their cell surface (**Fig 3A**). We inoculated both cell types with CHIKV-AF at a multiplicity of infection (MOI) of 1 or 10 and measured viral antigen (E2) expression by flow cytometry at 24 hpi. Whereas many (∼40 to 95%) fibroblasts were infected with CHIKV at either MOI, fewer (∼1 to 20%) endothelial cells were infected (**Fig 3B-C**). As an alternative approach, we performed experiments using blocking mAbs against MXRA8 [13]. Pretreating fibroblasts, but not endothelial cells, with anti-MXRA8 mAbs reduced CHIKV infection (**Fig 3D**).

**Figure 3.**
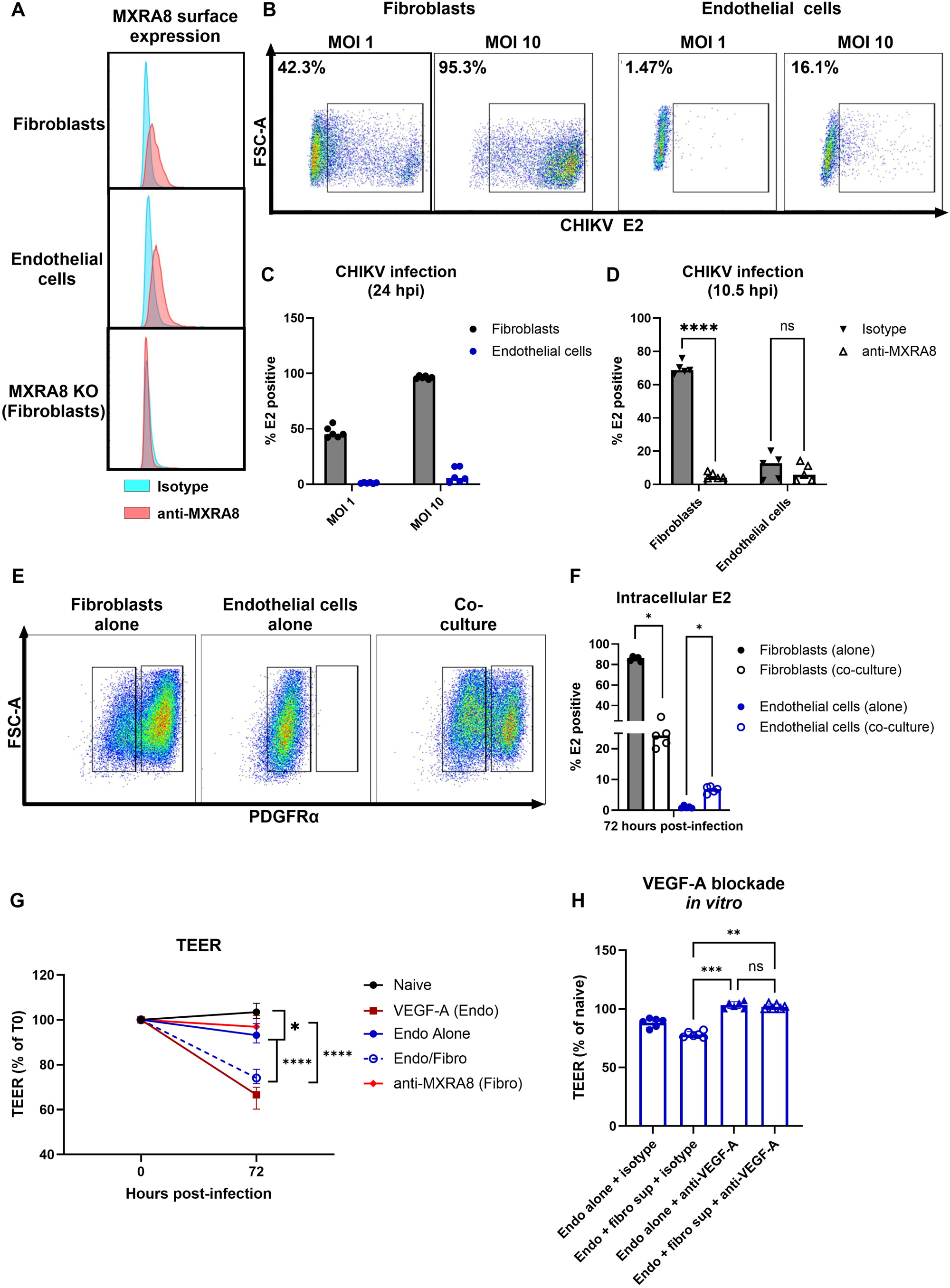
Fibroblast and endothelial cell infections with CHIKV. (**A**) NIH 3T3 fibroblasts or SVEC4-10 endothelial cell lines were sequentially incubated with hamster anti- MXRA8 antibodies (clones 1H1.F5 and 7F1.D8 [13]) and an AlexaFluor 647 conjugated anti-hamster IgG secondary antibody. MXRA8 surface expression was quantified by flow cytometry. (**B-C**) NIH 3T3 (*left panel*) or SVEC4-10 cells (*right panel*) were inoculated with CHIKV-AF at the indicated MOIs. At 24 hpi, cells were stained sequentially with an anti-CHIKV E2 mAb (CHK- 152, [68]) and an AlexaFluor 647-conjugated anti-mouse IgG secondary antibody and then processed by flow cytometry. (**D**) Fibroblasts or endothelial cells were pre-incubated with anti- MXRA8 antibodies or isotype control antibodies. Cells were infected at a MOI of 10, then collected at 10.5 hpi. (**E-F**) Fibroblasts or endothelial cells were plated alone or in a 1:1 co-culture, infected with CHIKV-AF at a MOI of 1, then collected and stained with anti-PDGFRα (**E**) or anti-CHIKV E2 mAbs (**F**). (**G**) Endothelial cells were plated in the top chamber of a transwell dish, and fibroblasts were plated in the bottom chamber. Cells in both chambers were infected with CHIKV-AF at a MOI of 1 and TEER was measured at 72 hpi. Where specified, VEGF-A was added to top chamber of the transwell, and anti-MXRA8 antibodies were added to the bottom chamber. (**H**) Endothelial cells were infected at a MOI of 1 with virus generated from infectious clones (endo alone) or from supernatants of CHIKV-infected fibroblasts collected at 72 hpi. Virus from infectious clones or from infected supernatants were diluted into PBS containing either anti-VEGF-A or isotype control antibodies. TEER was measured as in (**G**). Statistical analysis: (**D and F**) Mann-Whitney test, (**G-H**) Kruskal-Wallis ANOVA with Dunn’s post-test, *P\** < 0.05, \*\**P* < 0.01, \*\*\**P* < 0.001, P**** < 0.0001, ns, not significant.

Given that MXRA8 expression is similar in fibroblasts and endothelial cells, we reasoned that reduced endothelial cell infection efficiency was not dictated by differences in MXRA8- mediated entry. One possible explanation for reduced permissiveness to CHIKV infection is that endothelial cells express higher basal levels of antiviral host defense molecules [37, 38]. Accordingly, we measured expression of selected interferon (IFN)-stimulated genes (ISGs) with documented inhibitory activity against mosquito-transmitted RNA viruses [39–41] (*e.g*., *Ifit1* and *Oas1*) by qRT-PCR. Expression of *Oas1a* and *Ifit1* was increased in SVEC4-10 cells at baseline compared to 3T3 fibroblasts (**Fig S6A**). Similarly, we observed elevated basal expression of *Oas1a* in primary endothelial cells compared to fibroblasts derived from WT mice (**Fig S6B**). Moreover, blockade of IFN alpha and beta receptor subunit 1 (IFNAR1) with antibodies enhanced CHIKV-AF infection in endothelial cells (**Fig S6C**). Together, these data suggest that endothelial cells can be infected with CHIKV by a MXRA8-independent pathway, but the efficiency is likely restricted at a post-entry step by cellular host defense responses.

Because CHIKV infection in endothelial cell is reduced in *Pdgfra*-*Cre^ERT2^ Mxra8^fl/fl^* mice (**Fig 2F, H**) even though *Mxra8* expression is not altered (**Fig S5A**), we hypothesized that MXRA8-dependent fibroblast infection might facilitate endothelial cell infection. To begin to test this hypothesis, we inoculated 3T3 fibroblasts, SVEC4-10 endothelial cells, or co-cultures of the two cell types with CHIKV-AF (MOI of 1) and measured intracellular viral E2 antigen at 72 hpi. To distinguish fibroblasts and endothelial cells, we also stained cells for expression of PDGFRα. As expected, a majority of 3T3 cells (∼80%, **Fig 3E**, *left panel*) expressed PDGFRα, whereas SVEC4-10 cells did not (**Fig 3E**, *middle panel*). Compared to the isolated cultures, fibroblast infection was decreased in the co-culture system at 72 hpi (**Fig 3F**), suggesting that the presence of endothelial cells can reduce fibroblast infection efficiency with CHIKV. However, infection was increased (5-fold, *P* = 0.0079) in endothelial cells co-cultured with fibroblasts compared to those cultured alone (**Fig 3F**).

Endothelial cells have a critical role in maintaining vascular integrity [42], and loss of this function can lead to capillary leakage and tissue edema [43]. Given that deletion of MXRA8 from PDGFRα^+^ cells resulted in decreased endothelial cell infection and foot swelling, we speculated that CHIKV infection might alter endothelial cell barrier function, and this effect might be MXRA8-dependent. To begin to test this idea, we inoculated SVEC4-10 cells cultured alone in the top chamber of a transwell system or in the presence of 3T3 fibroblasts (in the bottom chamber) with CHIKV (MOI of 1) and measured trans-endothelial electrical resistance (TEER) at 72 hpi. As a positive control, we treated endothelial cells with vascular endothelial growth factor A (VEGF-A), which destabilizes adherens junctions and increases endothelial cell permeability [44]. SVEC4-10 endothelial cells cultured alone and infected with CHIKV had a small, but statistically significant, decrease in TEER (*P* = 0.04) compared to values just prior to infection (**Fig 3G**). However, endothelial cells co-cultured with fibroblasts had substantially reduced TEER after CHIKV infection compared to mock-infection (*P* < 0.0001) or CHIKV-infected endothelial cells cultured alone (*P* < 0.0001). This TEER change was reversed by treatment with blocking anti-MXRA8 mAbs (**Fig 3G**). These data support the idea that MXRA8-dependent CHIKV infection of fibroblasts influences endothelial cell permeability.

Fibroblasts are a known source of VEGF-A during homeostasis and in various pathological processes, including angiogenesis and cancer metastasis [45]. We hypothesized that CHIKV infection of fibroblasts might regulate endothelial cell permeability through secretion of VEGF. To test this idea, we inoculated endothelial cells at an MOI of 1 using either virus generated from infectious clones (endo alone) or virus from supernatants of CHIKV-infected fibroblasts at 72 hpi (fibro sup) along with 100 μg of anti-VEGF-A antibodies or isotype controls. We then measured TEER 24 h later. Endothelial cells inoculated with supernatants from CHIKV-infected fibroblasts showed reduced TEER that was comparable to what we observed in the co-culture model (**Fig 3G-H**). The reduction in TEER observed in the presence of fibroblast supernatant was rescued (*P* < 0.01) with anti-VEGF-A blockade compared to isotype antibody control-treatment (**Fig 3H**).

### Vascular permeability is increased during CHIKV infection in a MXRA8-dependent manner

Our co-culture studies suggest that CHIKV infection can reduce endothelial cell barrier integrity, and this phenotype is modulated by MXRA8-dependent fibroblast infection. To determine if vascular permeability in the foot is altered *in vivo* by CHIKV infection, we measured leakage into tissues of Evans Blue dye (C_34_H_24_Na_4_O_14_S_4_), which binds avidly to serum albumin [46]. Upon intravenous administration, the accumulation of Evans Blue dye in peripheral tissues reports microvascular leakage from circulation [47]. We inoculated five cohorts of mice with CHIKV-AF and at 1-, 2-, 3-, 5-, or 7 dpi administered 500 mg of Evans Blue dye via retro-orbital injection and allowed the dye to circulate for 45 min. Subsequently, mice were perfused extensively with PBS, and the foot ipsilateral to the site of CHIKV inoculation was harvested for dye extraction and measurement (**Fig 4A**). At 2 and 3 dpi, time points that correspond to the first peak of CHIKV-induced foot swelling in mice, we observed higher levels of Evans Blue in the ipsilateral feet of CHIKV-infected mice than in mock-infected controls (2 to 3-fold, *P* < 0.05) (**Fig 4B**), demonstrating that swelling correlates with vascular fluid leakage. We reasoned that if deletion of MXRA8 from PDGFRα-expressing cells abrogates foot swelling, it should reduce vascular leakage. To test this idea, we inoculated WT, *Mxra8*^Δ*8/*Δ*8*^ KO, tamoxifen-treated *Pdgfra*- *Cre^ERT2^ Mxra8^fl/fl^*, or tamoxifen-treated Cre*- Mxra8^fl/fl^* mice with CHIKV-AF, and at 3 dpi administered Evans Blue intravenously. Both global and PDGFRα-specific deletion of MXRA8 resulted in reductions (2 to 3-fold, *P* < 0.01) in Evans Blue dye translocation to the ipsilateral feet (**Fig 4C-E**).

**Figure 4.**
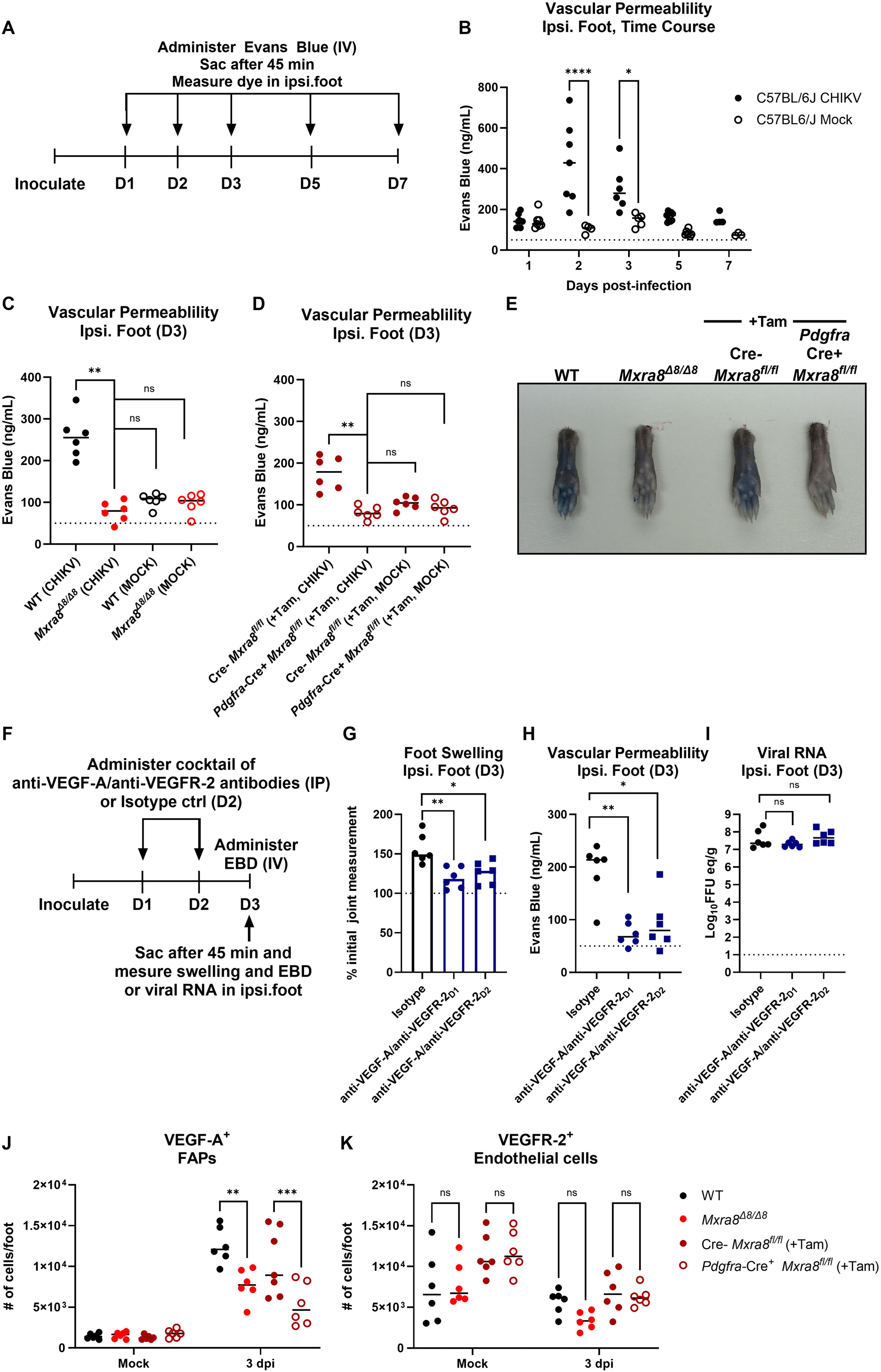
Vascular permeability in CHIKV-infected mice. (**A-B**) Cohorts of 4-week-old C57BL6/J WT mice (n = 6-7 mice per group, 2 experiments) were inoculated with 10^3^ FFU of CHIKV-AF in the left rear footpad or mock-infected. At 1, 2, 3, 5, or 7 dpi, Evans blue dye was administered intravenously via retroorbital injection to each cohort. After 45 min, mice were sacrificed, perfused with PBS, and the left rear foot was harvested. Evans blue dye was extracted from tissues and measured. (**C-E**) WT, *Mxra8*^Δ*8/*Δ*8*^ KO, tamoxifen (Tam)-treated *Pdgfra-Cre^ERT2^ Mxra8^fl/f^,* or Tam-treated Cre- mice were inoculated subcutaneously with CHIKV- AF in the left rear footpad. At 3 dpi, Evans blue dye was administered intravenously and extracted from left rear foot as in panel **B**. (**E**) Representative images of the ipsilateral feet from WT, *Mxra8*^Δ*8/*Δ*8*^*, Pdgfra-Cre^ERT2^ Mxra8^fl/fl^,* or *Cre-* mice at 45 min-after Evans blue dye administration. (**F-I**) Cohorts of 4-week-old C57BL6/J WT mice (n = 6-7 mice per group, 2 experiments) were inoculated with 10^3^ FFU of CHIKV-AF in the left rear footpad or mock- infected. (**F**) At 1 or 2 dpi, a 1:1 cocktail of anti-VEGF-A:anti-VEGFR-2 antibodies was administered via intraperitoneal injection. Foot swelling (**G**), vascular permeability (**H**), and viral RNA (I) was measured in the ipsilateral foot. (**J-K**) Cohorts of 4-week-old C57BL6/J WT, *Mxra8*^Δ*8/*Δ*8*^ KO*, Pdgfra-Cre^ERT2^ Mxra8^fl/fl^,* or *Cre-* mice (n = 6 mice per group, 2 experiments) were inoculated with 10^3^ FFU of CHIKV-AF in the left rear footpad or mock-infected. At 3 dpi, mice were administered brefeldin A via intraperitoneal injection 6 h prior to sacrifice. Ipsilateral feet were harvested and digested to release stromal cells. Cell suspensions were incubated with anti-CD45, anti-CD31, anti-CD29, anti-Sca-1, anti-VEGFR-2, and anti-VEGF-A antibodies, and the number and percentage of VEGF-A^+^ FAPs (**J**) or VEGFR-2^+^ endothelial cells (**K**) were determined using flow cytometry. Statistical analysis: (**B**) Two-way ANOVA with Šídák’s post-test (CHIKV-infected vs mock-infected) at each time point; (**C-D, G-I**) Mann-Whitney test; (**J-K**) Two- way ANOVA with Tukey’s post-test; \**P* < 0.05, \*\**P* < 0.01, \*\*\*\**P* < 0.0001; ns, not significant. Dotted horizontal lines on graphs indicate limit of detection for the assay.

Although VEGF-A signaling is a key regulator of vascular permeability [48, 49], its role in CHIKV pathogenesis is unknown. To determine if VEGF-A signaling contributes to the increase in permeability we observed in the ipsilateral feet during CHIKV infection (**Fig 4A**), we inoculated mice with CHIKV-AF, administered a 1:1 (500 μg total) cocktail of anti-VEGF-A:anti-VEGFR-2 mAbs or isotype control antibodies at 1 or 2 dpi via intraperitoneal injection and then measured foot swelling and Evans blue dye leakage into the tissue at 3 dpi (**Fig 4F**). VEGF-A signaling blockade at either day 1 or 2 dpi resulted in decreases in ipsilateral foot swelling (**Fig 4G**, 1.5 to 2-fold, *P* < 0.05) and vascular permeability (**Fig 4H**, 2-fold, *P* < 0.05) at 3 dpi. To determine if anti-VEGF-A signaling blockade affected virus infection, we measured CHIKV-AF viral RNA levels after administration of anti-VEGF-A:anti-VEGFR-2 mAb cocktail. Global levels of CHIKV-AF RNA in the foot were unaffected by treatment with anti-VEGF-A:anti-VEGFR-2 antibodies (**Fig 4I**). These data demonstrate that CHIKV-induced tissue edema and associated increases in microvascular leakage depend on VEGF-A signaling.

Given a role for VEGF-A signaling in CHIKV-induced tissue edema, we assessed how this pathway might be regulated during infection of WT and *Mxra8*^Δ*8/*Δ*8*^ KO mice. We harvested the tissue of ipsilateral feet of WT or *Mxra8*^Δ*8/*Δ*8*^ KO mice, and measured intracellular levels of VEGF-A in FAPs and endothelial cells at 3 dpi by flow cytometry. To block secretion of VEGF-A, we administered brefeldin A via intraperitoneal injection 6 h prior to sacrifice. The number of VEGF-A^+^ FAPs isolated from the ipsilateral feet of *Mxra8*^Δ*8/*Δ*8*^ mice was reduced compared to WT mice at 3 dpi (**Fig 4J**), a timepoint when we observed vascular leakage (**Fig 4B-E**). There also were fewer VEGF-A^+^ FAPs in the feet of tamoxifen-treated, CHIKV-infected, *Pdgfra*-*Cre^ERT2^ Mxra8^fl/fl^* mice compared to tamoxifen-treated, CHIKV-infected, Cre- *Mxra8^fl/fl^* controls (**Fig 4J**); this finding was not due to a MXRA8-dependent reduction in the total number of FAPs in the foot (**Fig S7A**). We measured VEGFR-2 expression on endothelial cells to determine if levels of the receptor for VEGF-A were altered during CHIKV infection in a MXRA8-dependent manner. However, differences in VEGFR-2 expression on endothelial cells were not detected between WT and *Mxra8*^Δ*8/*Δ*8*^ mice or *Pdgfra*-*Cre^ERT2^ Mxra8^fl/fl^* and Cre- *Mxra8^fl/fl^*mice either at baseline or during CHIKV infection at 3 dpi (**Fig 4K**). The total number of endothelial cells also did not differ between WT and *Mxra8*^Δ*8/*Δ*8*^ mice or Cre- and *Pdgfra*-*Cre^ERT2^ Mxra8^fl/fl^* mice (**Fig S7B**).

### The role of MXRA8 in CHIKV dissemination from infected feet

Although CHIKV can enter the bloodstream at early times (12 h) after infection via lymphatic transit of free virions [23] it remains unclear if virus also enters circulation from the underlying infected parenchymal tissues and what its contribution is for spread to distal tissues. Indeed, global [14] or PDGFRα- specific deletion of MXRA8 reduces CHIKV and MAYV dissemination to distal tissues (**Fig 2C- E, S3D-E**), which might reflect changes in vascular integrity and virus translocation (**Fig 4D**). We hypothesized that direct entry of CHIKV from the infected ipsilateral foot into circulation might coincide with vascular leakage and serve as a second pathway for dissemination.

To test this mechanism *in vivo*, we inoculated cohorts of WT or *Mxra8*^Δ*8/*Δ*8*^ KO mice with CHIKV-AF, and at 1, 2, or 3 dpi, injected into the ipsilateral foot 10^6^ FFU equivalents of the chimeric EILV-CHIKV, which contains the 5′ and 3′ untranslated regions, non-structural proteins, and subgenomic promoter of Eilat virus (EILV), an insect-specific alphavirus that is defective for replication in vertebrate cells [50], and the structural proteins (E1, 6 K, E2, E3 and capsid) of the Asian lineage CHIKV strain 99659 [51]. Like its parental virus, EILV-CHIKV only can replicate in insect cells and not in mouse or human cells. Forty-five minutes after injection, serum and the draining popliteal lymph nodes (DLN) were harvested for measurement of EILV RNA by qRT-PCR (**Fig 5A**). When EILV-CHIKV was administered at 1 dpi, EILV RNA levels were equivalent in the serum of WT or *Mxra8*^Δ*8/*Δ*8*^ KO mice. However, when EILV-CHIKV was given at 2 dpi, ∼10-fold lower (*P* < 0.001) levels of EILV *nsP1* RNA were detected in the serum of *Mxra8*^Δ*8/*Δ*8*^ KO than in WT mice (**Fig 5B**), and this difference was sustained when the EILV- CHIKV was injected at 3 dpi (∼20-fold, *P* < 0.01). When EILV-CHIKV was given to mock-infected mice, EILV *nsP1* RNA was undetectable in serum (**Fig 5B**), suggesting limited passive transit of EILV-CHIKV virions into the blood in the absence of active infection and inflammation. In comparison, EILV-CHIKV levels in the DLN were similar at 3 dpi in WT and *Mxra8*^Δ*8/*Δ*8*^ KO mice (**Fig 5C**), suggesting that the differences in viremia were not due to changes in efferent lymphatic transit in the absence of MXRA8. We did, however, observe a trend toward reduced lymphatic trafficking of EILV-CHIKV virions in both infected WT (*P* < 0.0001 at 2 dpi and *P* < 0.05 at 3 dpi) and *Mxra8*^Δ*8/*Δ*8*^ (*P* = 0.05 at 2 dpi) compared to their mock-infected counterparts.

**Figure 5.**
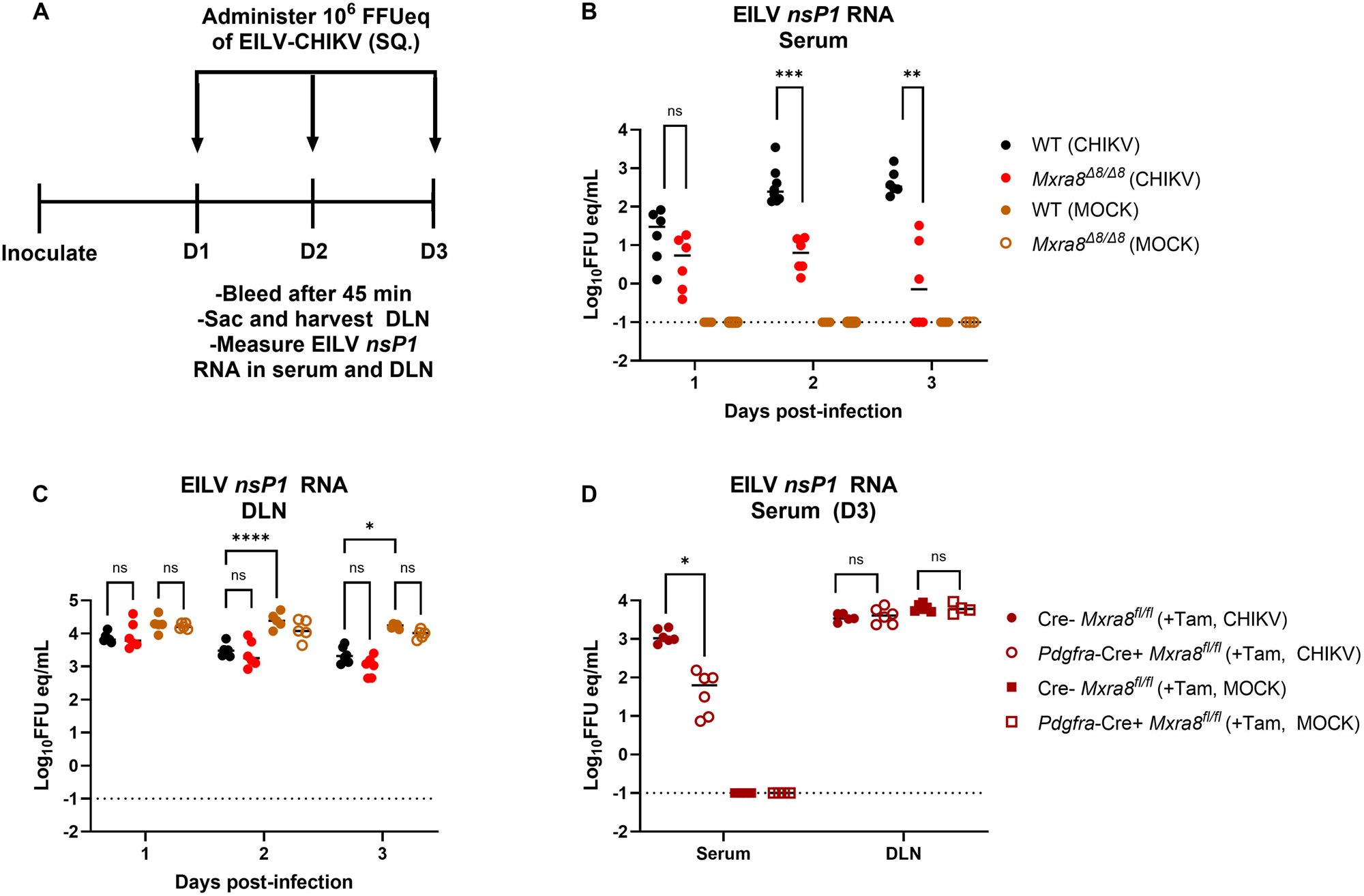
The effect of MXRA8-dependent infection on alphavirus translocation into circulation. (**A**) Experimental design. Cohorts of 4-week-old male C57BL6/J WT or *Mxra8*^Δ*8/*Δ*8*^ KO mice (n = 6-7 mice per group, 2 experiments) were inoculated with 10^3^ FFU of CHIKV-AF subcutaneously in the left rear footpad. Mock-infected animals were included as negative controls. At 1, 2, or 3 dpi, 10^6^ FFU equivalents of chimeric EILV-CHIKV (C-E3-E2-6K- E1 from CHIKV strain 99659) were injected subcutaneously in the left rear footpad. After 45 min, mice were bled, sacrificed, and draining popliteal lymph nodes (DLN) were collected. EILV non- structural protein 1 (*nsP1*) RNA was measured in serum (**B**) or DLN (**C**). (**D**) Tamoxifen (Tam)- treated *Pdgfra-Cre^ERT2^ Mxra8^fl/fl^,* or Tam-treated Cre- *Mxra8^fl/fl^* mice were inoculated subcutaneously with CHIKV-AF in the left rear footpad. At 3 dpi, 10^6^ FFU equivalents of EILV- CHIKV were injected subcutaneously in the left rear footpad. Mice were sacrificed, bled, and DLN was collected as in **B** and **C**. EILV non-structural protein 1 (*nsP1*) RNA was measured in serum or DLN. Bars indicate median values. Statistical analysis: (**B-D**) Mann-Whitney test; \**P* < 0.05, \*\**P* < 0.01, \*\*\**P* < 0.001, \*\*\*\**P* < 0.0001. Dotted horizontal lines on graphs indicate limit of detection for the assay.

To determine if alphavirus particle trafficking from the ipsilateral foot to the circulation specifically depends on MXRA8 expression in PDGFRα^+^ cells, we repeated the EILV-CHIKV injection experiment in CHIKV-AF-infected, Cre- and *Pdgfra*-*Cre^ERT2^ Mxra8^fl/fl^* mice. EILV RNA levels in serum were lower (10-fold, *P* < 0.01) in *Pdgfra*-*Cre^ERT2^ Mxra8^fl/fl^* mice than in Cre- littermates (**Fig 5D**) but were unchanged in the DLN (**Fig 5D**). Together, these data suggest that alphavirus virions can transit into the bloodstream from underlying infected peripheral tissues at time points that correspond to enhanced vascular leakage, and that this dissemination mode requires MXRA8 expression in PDGFRα^+^ cells.

## DISCUSSION

In this study, in musculoskeletal tissues, we observed that CHIKV infection of PDGFRα^+^mesenchymal cells but not endothelial cells depends on expression of the entry receptor, MXRA8. Nonetheless, endothelial cell infection efficiency was enhanced by MXRA8-dependent CHIKV infection of PDGFRα^+^ mesenchymal cells, which together promoted vascular leakage, tissue edema, and hematogenous virus dissemination. Together, with prior studies, our experiments elucidate the early dynamics of CHIKV tropism and dissemination. At early times post-infection (18 to 24 hpi) CHIKV enters the bloodstream through efferent lymphatic transit, and this pathway is dependent on inflammatory monocytes [23]. However, soon afterwards (2 to 3 dpi), stromal VEGF-A production, likely by infected fibroblasts, decreases endothelial cell barrier integrity, which enables CHIKV to directly enter the bloodstream from underlying infected parenchymal tissues.

In vasculature, PDGFRα^+^ cells reside in the *tunica adventitia*, the outermost layer of blood vessels [52]. Given their proximity, one possible explanation for the role of PDGFRα^+^ cells in promoting CHIKV infection of endothelial cells is increased local production of virus and direct spread. In our co-culture model, the presence of fibroblasts enhanced CHIKV infection of endothelial cells in a MXRA8-dependent manner. Pericytes, another perivascular cell type, have been shown to play an analogous role in modulating endothelial barrier integrity and infection during Japanese encephalitis [53], Andes [54], and Zika [55] virus infections. Although we observed that infection of pericytes in skeletal muscle occurs through a MXRA8-independent mechanism, we cannot rule out a contribution of these cells to CHIKV-induced vascular leakage. Indeed, in an *in vitro* blood-brain barrier model, pericyte infection by Sindbis virus, a distantly related alphavirus, decreased barrier integrity of brain microvascular endothelial-like cells through infection via its entry receptor, protocadherin-10 (PCDH10) [56].

Sustained disruption or damage to endothelial barrier function after virus infections can lead to edema, shock, and hemorrhagic syndromes [57]. Given the importance of maintaining barrier integrity, there may be evolutionary adaptations to limit endothelial cell infection by viruses. Indeed, human umbilical vein endothelial cells (HUVECs) display transcriptional signatures of type I IFN signaling and phosphorylated STAT1 in the absence of inflammatory stimuli [37]. We observed elevated basal expression of the ISGs *Oas1a* and *Ifit1* in transformed and primary endothelial cells compared to fibroblasts. Consistent with this data, endothelial cells are inherently resistant to infection by many viruses [37, 58–61]. In human cardiac microvascular endothelial cells, CHIKV infection is rescued upon treatment with ruxolitinib, a JAK/STAT inhibitor, suggesting that restriction of infection is, in part, at the level of cell-intrinsic immunity [38]. In a co-culture model, basal type I IFN responses in HUVECs protect neighboring epithelial cells from SARS-CoV-2 infection [37]. In our co-culture model, although the presence of fibroblasts enhanced endothelial cell infection, fibroblast infection was reduced by the presence of endothelial cells. It will be important to determine the specific mechanisms by which arthritogenic alphaviruses are controlled in endothelial cells and the impact this has on neighboring cells in the tissue.

The VEGF family of growth factors is comprised of four different isoforms (VEGF-A, -B, - C, -D) and placental growth factor [62]. As a potent inducer of vascular permeability and a specific endothelial cell mitogen, VEGF-A binds to VEGF receptor 2 (VEGFR-2) on endothelial cells and regulates angiogenesis [62]. Our experiments showed that blockade of VEGF-A signaling in WT mice reduced CHIKV-induced foot swelling and vascular permeability. Given that these phenotypes are also reduced in *Mxra8*^Δ*8/*Δ*8*^ KO and *Pdgfra*-*Cre^ERT2^ Mxra8^fl/fl^* mice, we hypothesized that VEGF-A expression might be reduced in FAPs. In wound repair models, fibroblasts are a key source of VEGF-A [63]. During CHIKV infection in mice, we observed fewer VEGF-A-expressing FAPs in both *Mxra8*^Δ*8/*Δ*8*^ KO and *Pdgfra*-*Cre^ERT2^ Mxra8^fl/fl^* mice. Collectively, our data suggest that CHIKV induced MXRA8-dependent infection of FAPs may produce VEGF- A and trigger endothelial cell changes in vascular permeability. More investigation is warranted to define the upstream signals that regulate VEGF-A expression in the context of CHIKV infection and how this results in tissue edema.

MXRA8-dependent vascular leakage promoted hematogenous dissemination of non-replicating EILV-CHIKV virions starting at 2 dpi, which contrasts with the earlier phase of viremia (*i.e*., 18 hpi) that originates from lymphatic transit [23]. CHIKV has likely evolved multiple routes of dissemination to the bloodstream, possibly to evade clearance by scavenger receptors (*e.g*., MARCO [64]) in lymphatic tissues or to increase the magnitude of viremia and likelihood of transmission back to mosquitoes. Given that viremia also peaks between 2 and 3 dpi during CHIKV infection of mice and humans [64], hematogenous dissemination from the initial site of musculoskeletal tissue infection may be more efficient than lymphatic spread.

We acknowledge several limitations of our study. (a) We did not identify the signal responsible for initiating VEGF-A expression in FAPs during CHIKV infection or the signaling cascade in endothelial cells that regulates vascular permeability. While we show that FAPs produce VEGF-A in a MXRA8-dependent manner and that MXRA8-dependent infection is required for foot swelling, we did not directly demonstrate that VEGF-A produced by FAPs is required. Conditional deletions of *Vegfa* in FAPs could begin to address the question. (b) We did not assess the contribution or mechanistic role of other CHIKV-permissive cell types (*e.g*., skeletal muscle cells, monocytes) to vascular permeability. Indeed, monocytes promote CHIKV-induced foot swelling in mice [65, 66] although their linkage to VEGF-A or other signaling pathways that affect vascular permeability remains unknown. (c) We did not identify the MXRA8-independent pathway of CHIKV infection in the foot. Additional receptor discovery campaigns are needed to identify alternate pathways for alphavirus entry into cells.

In summary, CHIKV replicates in multiple cell types in the skin and underlying soft tissues to achieve efficient dissemination to the bloodstream. Our study provides evidence that MXRA8-dependent infection of PDGFRα^+^ mesenchymal cells and interactions with endothelial cells are critical for regulating CHIKV-induced foot swelling. We show that foot swelling is a consequence of CHIKV-induced increases in vascular permeability, which depends on MXRA8 expression and VEGF signaling. Finally, we demonstrate that these changes in vascular permeability enable a direct route for alphavirus virions to enter circulation.

## METHODS

### Cells and viruses

BHK21 cells (CCL-21, ATCC) Vero cells (CCL-81, ATCC), NIH 3T3 fibroblasts (CRL-1658, ATCC), and SVEC4-10 murine endothelial cells (CRL-2181, ATCC) were cultured in Dulbecco’s Modified Eagle Medium (DMEM, Gibco) supplemented with 5% heat-inactivated fetal bovine serum (FBS) at 37°C with 5% CO_2_. CHIKV AF15561, CHIKV AF15561- Venus (a gift of T. Morrison, University of Colorado School of Medicine), CHIKV LR2006 OPY1, and MAYV BEH04 were generated from infectious cDNA clones as described previously [14, 23]. Briefly, viral RNA was *in vitro* transcribed from cDNA template using mMESSAGE mMACHINE SP6 Transcription Kit (ThermoFisher, AM1340). Viral RNA was electroporated into BHK21 cells, and supernatants containing infectious virus were collected 48 h later. Virus titer was measured by plaque assay on Vero cells. Viruses were then passaged once (CHIKV- AF15561) or twice (CHIKV-AF15561-Venus and MAYV) on BHK21 cells. EILV-CHIKV [67] was obtained from Dr. Scott Weaver and the World Reference Center for Emerging Viruses and Arboviruses (University of Texas Medical Branch) and passaged once in *Aedes albopictus* C6/36 cells (CRL-1660, ATCC).

### Mice

All mouse experiments were performed according to the *Guide for the Care and Use of Laboratory Animals* of the National Institutes of Health after approval by the Institutional Animal Care and Use Committee at the Washington University School Medicine. All infections in mice were performed under anesthesia with ketamine hydrochloride (80 mg/kg) and xylazine (15 mg/kg). Four-week-old WT C57BL/6J mice were purchased from Jackson Laboratories (000664). Conditional deletion of *Mxra8* was generated by introducing *loxP* sites flanking the second and last exons of the *Mxra8* coding region (**Fig S2A**). Two sgRNAs were selected based on a low off-target profile and distance to the target site: sgRNA-1: 5’- AGCTGTTCTAGCATGACACTNGG-3’ and sgRNA-2: 5’ TAGCAGTGTGATCTCTGATGNGG-3’. The two gRNAs were generated by Integrated DNA Technologies. Guide RNAs, CAS9 protein, and the oligo donor sequences were complexed and electroporated into C57BL/6J zygotes. After electroporation, founder lines were confirmed by genotyping and next-generation sequencing using the Illumina 2x250 MiSeq platform. Founders were backcrossed for two generations to obtain homozygous floxed (*Mxra8*^fl/fl^) mice. *Mxra8*^fl/fl^ mice were then crossed to *Pdgfra*-*Cre^ERT2+^*(018280, Jackson Laboratories) or *Vav1*-Cre (035670, Jackson Laboratories). Littermate Cre-negative, *Mxra8*^fl/fl^ mice were used as controls. Tamoxifen (2 mg/mouse, T5648, MilliporeSigma) was dissolved in corn oil and administered to *Pdgfra*-*Cre^ERT2+^ Mxra8*^fl/fl^ mice every other day for 5 days to induce Cre-recombinase expression. *Mxra8*^fl/fl^ mice were generated with the aid of the Genome Engineering and iPSC center and Department of Pathology Micro-Injection Core (Washington University School of Medicine). *Mxra8*^Δ*8/*Δ*8*^ global KO mice were described previously [14].

### *In vitro* infections

SVEC4-10 endothelial cells (2 x 10^5^ cells) or NIH 3T3 fibroblasts (2 x 10^5^ cells) were seeded into 96-well plates. Virus was diluted into 100 μL of DMEM containing 2% FBS, added to cell monolayers, and incubated at 37°C with 5% CO_2_ for specified time points. Cells were collected with TrypLE (Gibco), fixed in 4% paraformaldehyde (PFA), permeabilized in Hank’s Balanced Salt Solution (HBSS) buffer containing 0.1% saponin (w/v)and 2% FBS for 15 min at room temperature. Cells were incubated with mouse mAbs (CHK-152 [68]) against the CHIKV E2 protein for 30 min at 4°C, washed twice with FACS buffer (2% FBS and 2 mM Ethylenediaminetetraacetic acid [EDTA] in PBS), then incubated with anti-mouse secondary antibodies conjugated to AlexaFluor 647. Cells were washed again twice with FACS buffer, then stored in 0.1% PFA in 2% FBS until analysis by flow cytometry. For transwell infections, endothelial cells (2 x 10^5^ cells) were plated in the top chamber, and fibroblasts (2 x 10^5^ cells) were plated in the bottom chamber. Virus was diluted into 300 μL of DMEM containing 2% FBS and added to either chamber. Cells were then assessed for intracellular E2 expression

### Mouse infections

Four-week-old male or female WT C57BL6/J, *Mxra8*^Δ*8/*Δ*8*^ KO*, Pdgfra- Cre^ERT2+^ Mxra8*^fl/fl^, or *Vav1*-Cre *Mxra8*^fl/fl^ mice were inoculated in the footpad with 10^3^ FFU of CHIKV AF15561, 10^3^ FFU of MAYV, or 10^5^ FFU of CHIKV-AF15561-Venus diluted in Dulbecco’s phosphate buffered saline (DPBS, Gibco). At 3 dpi, swelling of the ipsilateral foot (width x height) was measured using digital calipers. Mice were sacrificed with a ketamine/xylazine overdose per Washington University School of Medicine Institutional Animal Care and Use Committee guidelines and perfused with 20 mL of DPBS. Tissues were harvested and flash frozen on dry ice prior to storage at −80°C.

### Quantification of viral RNA or infectious virus particles

Viral RNA was extracted from homogenized tissues or infected cell lines and quantified as described previously [14, 23]. Primer and probe sets for CHIKV AF15561 *E2*, MAYV *E2*, and EILV *nsP1* were previously reported [23].

### TEER assays

Endothelial cell TEER was measured using a Millicell® ERS 3.0 Digital Voltohmmeter (MilliporeSigma) just prior to infection and at 3 days after CHIKV-AF infection. Ohm values obtained from blank wells were subtracted from values obtained from treatment wells to account for background. Values are represented as a percentage of the TEER measured at day 0.

### Vascular permeability assays

At 1, 2, 3, 5, or 7 dpi, mice were anesthetized with ketamine/xylazine cocktail. Subsequently, 100 μL of Evans Blue dye (5 mg/mL in DPBS) was administered intravenously via retro-orbital vein injection. Mice were sacrificed at 45 min post-injection, perfused with 20 mL of DPBS, and ipsilateral feet were collected. Tissue was cut into small pieces and incubated in 200 μL of formamide (Cat # F9037, MilliporeSigma) at 65°C for 48 h to extract the dye. Optical density (OD) was measured using a spectrophotometer with an excitation wavelength of 620 nm. Measurements are reported as ng/mL and were calculated using a standard curve. For virus trafficking experiments, at 1, 2, or 3 dpi, mice were anesthetized with ketamine/zylazine, and then 10^6^ genome equivalents of EILV-CHIKV were injected subcutaneously into the ipsilateral foot. Mock-infected mice were used as negative controls. At 45 min post-injection, mice were bled, sacrificed, and the draining popliteal lymph node was collected. Tissue was homogenized, and EILV-CHIKV RNA was quantified by qRT-PCR as described previously [23].

### Anti-VEGF-A treatments

Cohorts of four-week-old male WT C57BL6/J mice were inoculated in the footpad with 10^3^ FFU of CHIKV AF15561. At 1 or 2 dpi, mice were administered a 1:1 cocktail of anti-VEGF-A (BioXCell, cat # BE0399) and anti-VEGFR2 (BioXCell cat # BE0060) antibodies diluted in PBS (Gibco) or 500 ug of isotype control antibodies (250 ug BioXCell, cat # BE0089 and 250 ug BioXCell cat # BE0088) via intraperitoneal (IP) injection. Foot swelling was measured using digital calipers. Vascular permeability was measured using the Evans blue dye assay described above in the “**Vascular permeability assays**” section. Viral RNA was quantified as indicated in “**Quantification of viral RNA or infectious virus particles”** section.

### Flow cytometry

Ipsilateral feet were harvested at 3 dpi and incubated with 5 mg/mL of Collagenase IV (Gibco), 1 μg/mL of LiberaseTL (Roche), and 10 µg/mL of DNAse (Roche) diluted in RPMI (Gibco) for 1 hour at 37°C. Subsequently, tissue was passed through a 70-µm cell strainer and centrifuged at 1500 RPM for 5 min. Cells then were incubated with antibodies against CD45 (clone 30-F11, Biolegend, Cat # 103133), CD11b (clone M1/70, Biolegend, Cat # 101256), Ly6C (clone HK 1.4, Biolegend, Cat # 128016), CD31 (clone 390, Biolegend, Cat # 102430), CD29 (clone HMβ1-1, Biolegend, Cat # 102218), anti-VEGFR-2 (clone Avas12, Biolegend, Cat # 136406), CD146 (clone ME-9F1, Biolegend, Cat # 134714), eFluor 506 fixable viability dye (Invitrogen, Cat # 65086614), and TruStain FcX^tm^ anti-mouse CD16/32 (clone 93, Biolegend, Cat # 101320). Cells were fixed in 4% paraformaldehyde (PFA) and processed on a Cytek Aurora flow cytometer.

For assessment of intracellular VEGF-A expression, mice were inoculated with CHIKV as indicated in the “**Mouse Infections**” section. Six hours prior to sacrifice, mice were administered brefeldin A (MedChem, 250 ug/mouse in 200 uL PBS) via intraperitoneal injection. Ipsilateral feet were harvested, processed, and stained with the antibodies listed above. After initial surface staining, cells from foot tissues were fixed with 100 μL of Fix and Perm Medium A (FIX & PERM™ Cell Permeabilization Kit, Invitrogen) then permeabilized with 100 μL of Medium B according to manufacturer’s protocol. Cells were then incubated with anti-VEGF-A antibodies (Cat# NB100-664AF594 Novus Biologicals). Cells were washed twice with 2% BSA containing 2 mM EDTA in PBS then stored in 0.1% paraformaldehyde in PBS at 4°C. Analysis was performed on a Cytek Aurora flow cytometer and analyzed using FlowJo software, (FlowJo, LLC, version 10.8.1).

### RNA sequencing analysis

Analysis was carried out on previously published single cell RNA sequencing datasets (GEO accession GSE300299) [36] obtained from the mouse foot after CHIKV infection. Only barcodes carrying an original author-assigned cell type annotation were retained (n = 75,547), and re-clustering or re-annotation was not done; cell numbers per annotated population and sample recapitulate those reported in the original study. The three CHIKV viral transcripts present in the deposited matrix were removed prior to normalization, so that per-cell library sizes were calculated over host transcripts only. Counts were normalized to total counts per cell, scaled by a factor of 10,000 and natural-log transformed (NormalizeData, Seurat v5.4.0; [69]). For each gene and cell type, the percentage of cells in which the transcript was detected (>=1 UMI) and the mean log-normalized expression were computed, both pooled across all cells and separately for each of the six animals. All statements comparing groups are based on per-animal values.

### Statistical analysis

Statistical analyses were carried out using GraphPad Prism v 10.1.0. Mann-Whitney, Kruskal-Wallis ANOVA with multiple comparisons analysis, or 2-way ANOVA tests were used to determine statistical significance between experimental groups. A *P* value of < 0.05 was deemed significant.

## Data availability

All data supporting the findings of this study are available within the paper and its Supplementary data. This paper does not include original code. Any additional information or reagents related to the study are available from the corresponding author upon request.

## Acknowledgements

This study was supported by the National Institutes of Health (R01 AI143673 to M.S.D. and R01 AI183590 to D.J.L.) and The Hanna H. Gray Fellows Program of the Howard Hughes Medical Institute (to A.C.H.).

## Author contributions

A.C.H. and M.S.D. designed studies and wrote the initial draft, with all other authors providing editorial comments. A.C.H performed *in vitro* infections, mouse infections, tissue harvesting, mouse genotyping, RT-qPCR, flow cytometry, Evans blue permeability assays, virus particle trafficking assays, and plaque assays. Y.A. performed bioinformatics analysis on published single-cell RNA sequencing datasets [36]. D.J.L. and M.S.D. supervised studies and provided resources for the experiments.

## SUPPLEMENTAL FIGURE LEGENDS

**Figure S1.**
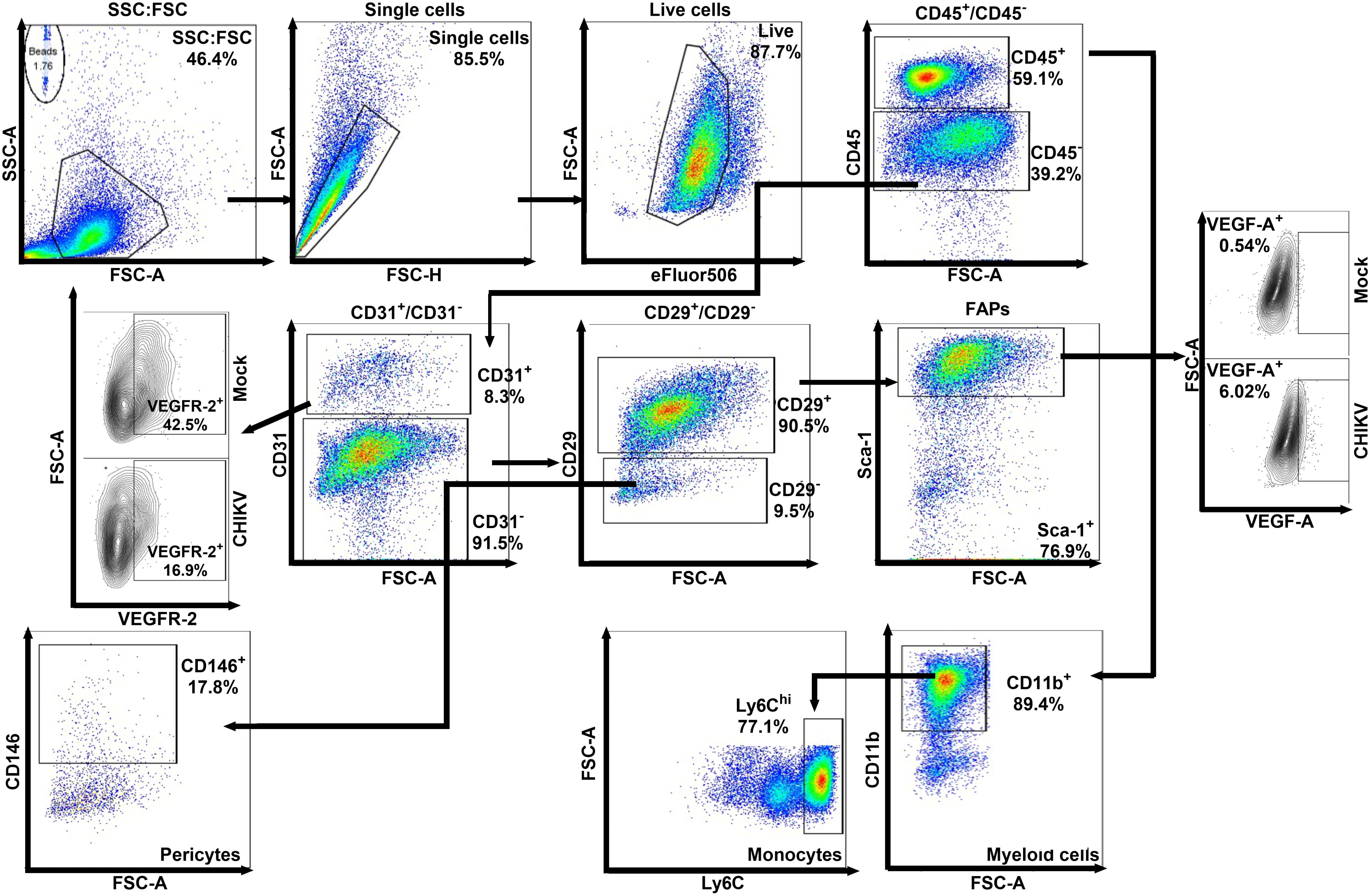
Flow cytometry gating scheme for CHIKV-infected cells. Non-hematopoietic cells were defined as live, single, CD45^-^ cells, and hematopoietic cells were defined as live, single, CD45^+^ cells. Both cell compartments were analyzed for YFP expression after CHIKV-AF15561-Venus infection. Naive cells were used to set gates. Monocytes: live, single, CD45^+^, CD11b^+^, Ly6C^hi^ cells; endothelial cells: live, single, CD45^-^, CD31^+^ cells; FAPs: live, single, CD31^-,^ CD29^+^, Sca-1^+^ cells; and pericytes: live, single, CD45^-^, CD31^-^, CD29^-^, CD146^+^ cells. FAPs were analyzed for intracellular VEGF-A expression and endothelial cells were analyzed for VEGFR-2 expression.

**Figure S2.**
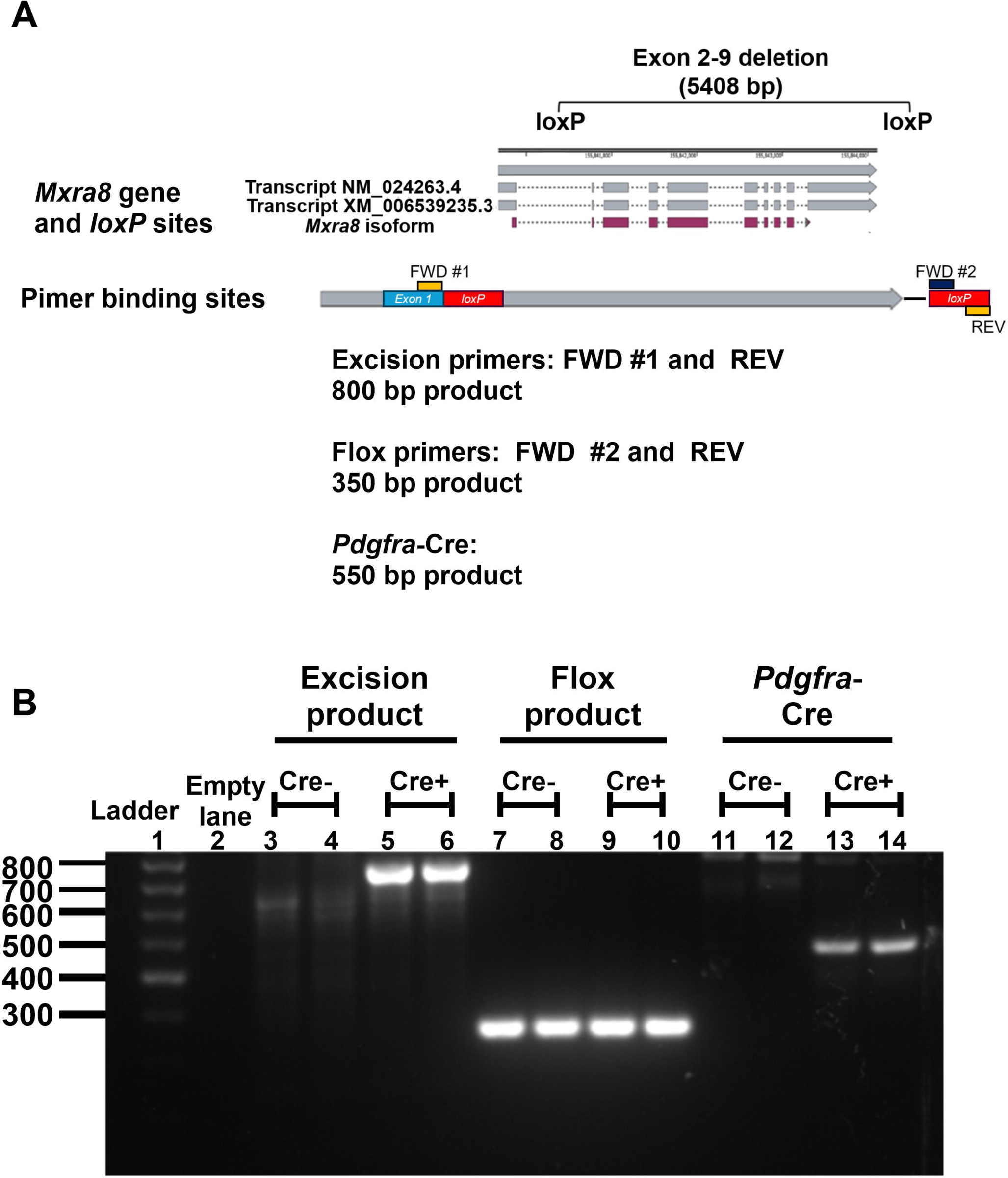
Genotyping of *Pdgfra-Cre^ERT2^ Mxra8^fl/fl^* mice. (**A**) Location of the *loxP* sites flanking exons 2-9 of the mouse *Mxra8* gene (*top panel*) and location of the primer binding sites for primers used to validate *Pdgfra-Cre^ERT2^ Mxra8^fl/fl^* mice. (**B**) Animals were administered tamoxifen (Tam) as indicated in the Methods. Tail sections were clipped from two separate *Pdgfra-Cre^ERT2+^*or Cre*-* mice and digested in lysis buffer. Genomic DNA was isolated and used as a template in three separate PCR reactions. Excised and floxed regions of the *Mxra8* gene were amplified by PCR using the following primers: Recombination product (lanes 3-6), Forward 5’ GTCCTGCTGTGGAAACTGCTGC 3’ and Reverse 5’ CTTGATTCCCAGCTGACTCGCC; Floxed product (lanes 7-10) Forward 5’ GTCCTGCTATGGAAACTGCTAC 3’ and Reverse 5’ CTTGATTCCCAGCTGACTCGCC*. Pdgfra-Cre^ERT2^*(lanes 11-14) was amplified using the following primers: Forward 5’ ATCGCATTCCTTGCAAAAGT 3’ and Reverse 5’ AGGCCCACAGAACATGGAC 3’.

**Figure S3.**
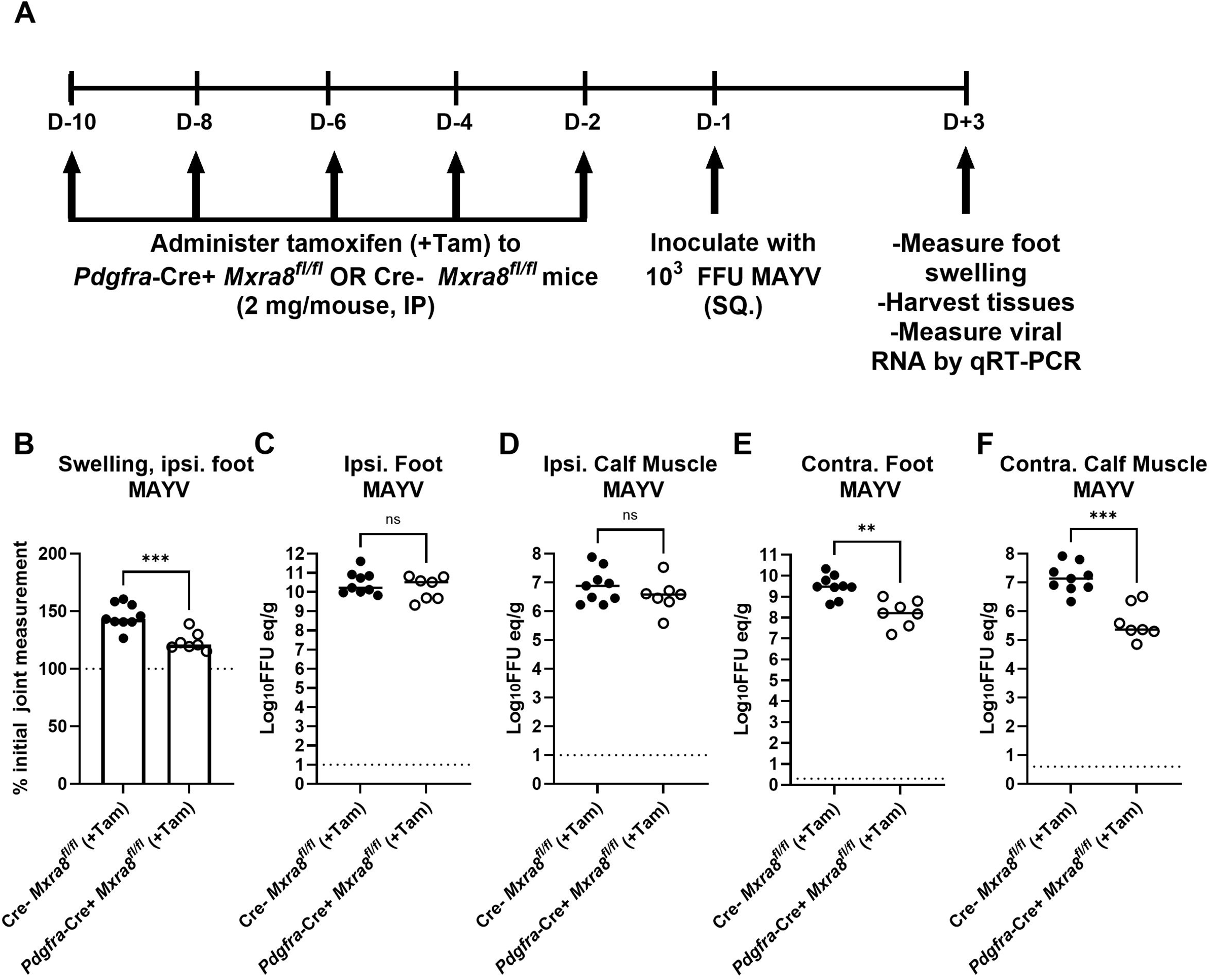
MAYV infection of *Pdgfra-Cre^ERT2^ Mxra8^fl/fl^* mice. (**A**) Three-week-old male and female *Pdgfra-Cre^ERT2^ Mxra8^fl/fl^* mice were administered tamoxifen (Tam) every other day for 5 days via intraperitoneal injection. Subsequently, *Pdgfra-Cre^ERT2^*^+^ or Cre- *Mxra8^fl/fl^* mice (n = 7-9 mice per group, 2 experiments) were inoculated with 10^3^ FFU of MAYV via subcutaneous footpad injection. (**B**) At 3 dpi, footpad swelling was measured. Mice were then sacrificed, perfused with PBS, and ipsilateral foot (**C**), ipsilateral calf muscle (**D**), contralateral foot (**E**), and contralateral calf muscle (**F**) were harvested. Viral RNA in each tissue was measured by qRT-PCR. Statistical analysis: (**B-F**) Mann-Whitney test; *P\** < 0.05, \*\**P* < 0.01, \*\*\**P* < 0.001, ns, not significant. Dotted horizontal lines on graphs indicate limit of detection for the assay.

**Figure S4.**
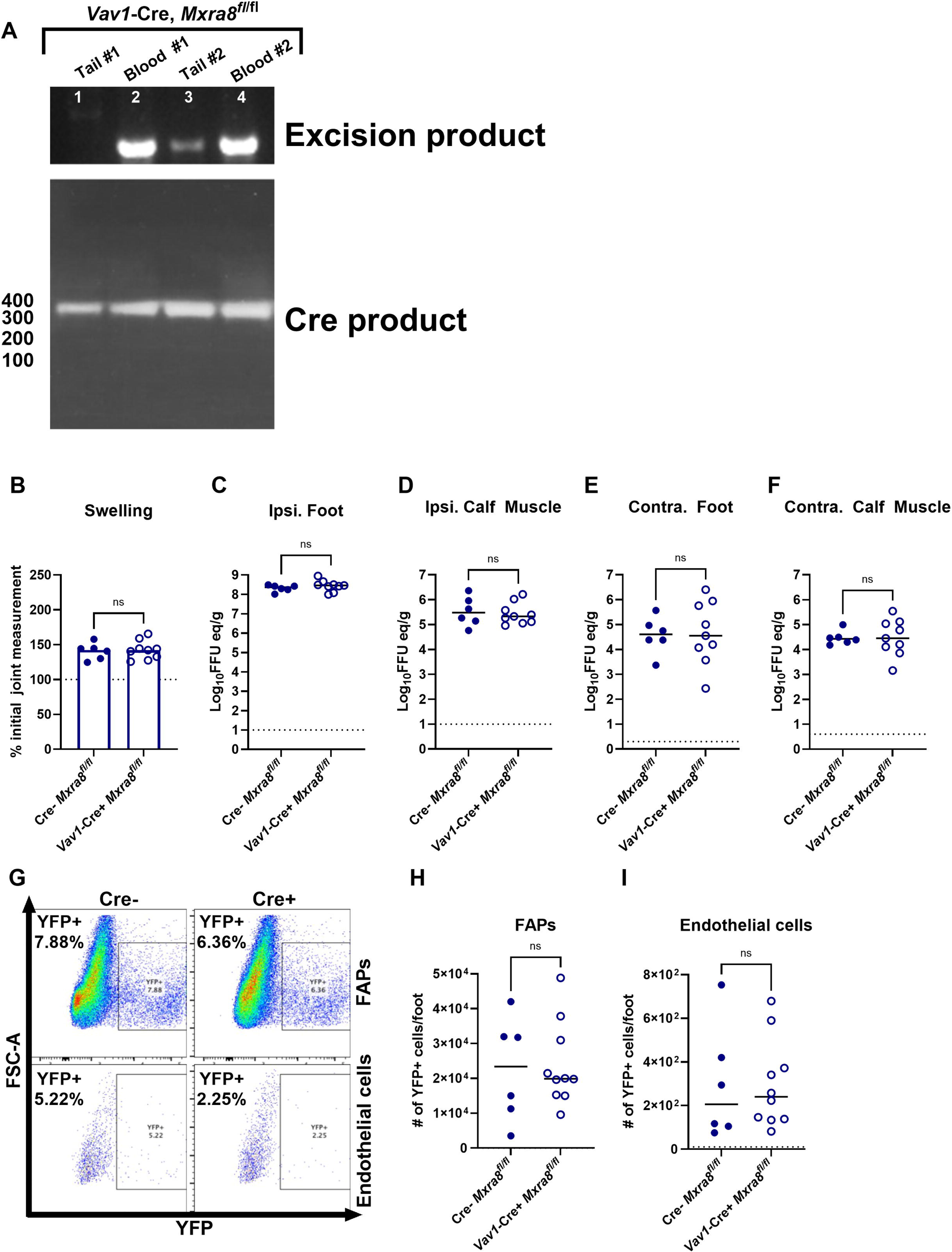
CHIKV infection of *Vav1-Cre Mxra8^fl/fl^* with CHIKV-AF or CHIKV-AF- Venus. (**A**) Tail sections were clipped or blood was collected from two separate *Vav1-Cre Mxra8^fl/fl^* mice. Tails or whole blood were digested in lysis buffer. Genomic DNA was isolated and used as a template in two separate PCR reactions to amplify products of *Mxra8* gene excision (*top panel*) or *Vav1*-Cre (*bottom panel*). (**B-F**) Four-week-old male and female *Vav1-Cre Mxra8^fl/fl^* mice (n = 6-10 mice per group, 2 experiments) were inoculated with 10^3^ FFU of CHIKV- AF via subcutaneous footpad injection. At 3 dpi, footpad swelling (**B**) was measured. Mice were then sacrificed, perfused with PBS, and the ipsilateral foot (**C**), ipsilateral calf muscle (**D**), contralateral foot (**E**), and contralateral calf muscle (**F**) were harvested. Viral RNA in each tissue was measured. (**G-I**). Cre- or *Vav1-*Cre *Mxra8^fl/fl^* mice were inoculated with 10^5^ FFU of CHIKV- AF-Venus in the footpad. At 2 dpi, ipsilateral feet were collected and processed as in Fig 1 **and 2**. The percentage (**G**) and number of FAPs (**H**) or endothelial cells (**I**) expressing YFP were quantified by flow cytometry. Statistical analysis: (**B-I**) Mann-Whitney test; ns, not significant. Dotted horizontal lines on graphs indicate limit of detection for the assay.

**Figure S5.**
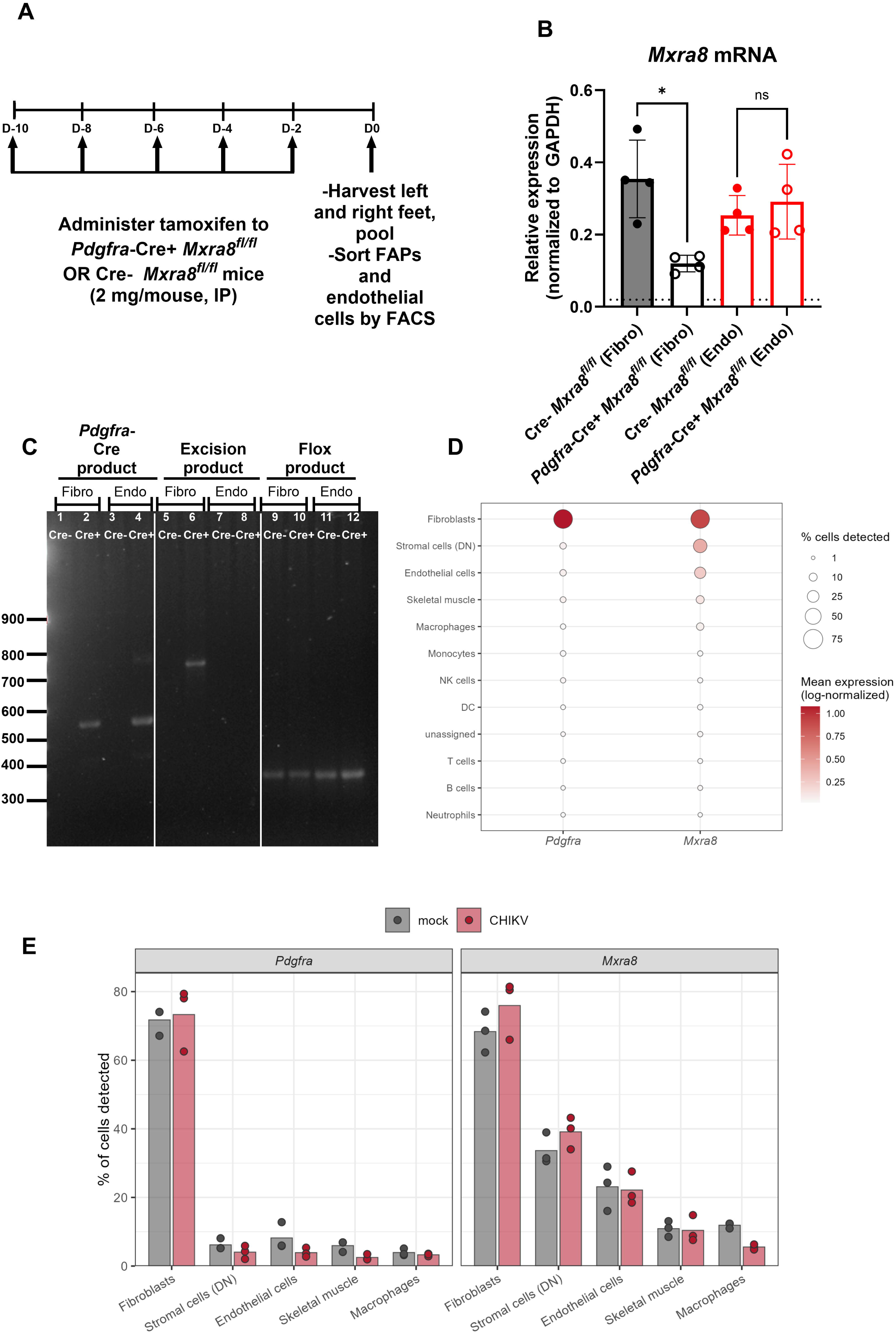
MXRA8 expression in fibroblasts and endothelial cells isolated from the feet of *Pdgfra-Cre^ERT2^ Mxra8^fl/fl^* mice. (**A**) Three-week-old male and female Cre- or *Pdgfra- Cre^ERT2^ Mxra8^fl/fl^*mice were administered tamoxifen (Tam) every other day for 5 days via intraperitoneal injection. Both feet were harvested and processed as in **Fig 1, 2, and S3**. Cells were pooled from 8 Cre+ or 8 Cre- mice, and the experiment was repeated four times. CD45^-^ cells were sorted into the CD31^+^ endothelial cells and CD31^-^ CD29^+^ fibroblast fractions. (**B**) Cellular mRNA was extracted from 10^5^ endothelial cells or fibroblasts to measure *Mxra8* mRNA levels by qRT-PCR and was normalized to *Gapdh* mRNA levels. (**C**) In a separate experiment, DNA was extracted from 10^5^ endothelial cells or fibroblasts and used a template to amplify regions of the genome corresponding to products of Cre-recombinase-mediated excision. (**D-E**) Processed single-cell RNA-seq data (count matrix and cell metadata) were downloaded from GEO accession GSE300299 [36]. (n = 3 per group; [70]). Statistical analysis: Mann-Whitney test; \**P* < 0.05. Dotted line indicates limit of detection for the assay.

**Figure S6.**
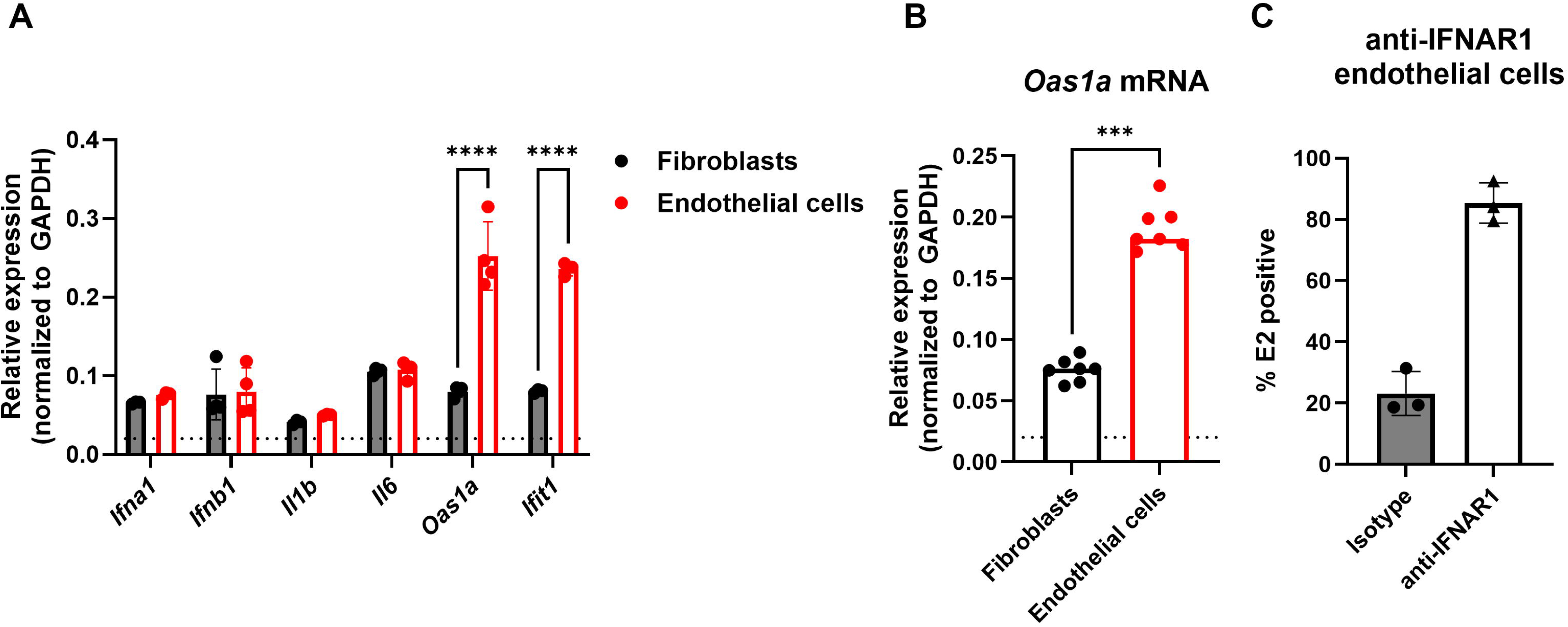
ISG expression and IFNAR1 blockade. (**A**) RNA was extracted from 2 x 10^5^ NIH 3T3 fibroblasts or SVEC4-10 cells and used as template to measure mRNA levels of a panel of genes (*Ifna*, *Ifnb*, *Il1b*, *Il6*, *Oas1a*, and *Ifit1*) by qRT-PCR. *Gapdh* levels were used to normalize between cell types. (**B**) Primary fibroblasts and endothelial cells were isolated, pooled, and sorted from both feet of WT mice (n = 7), and levels of *Oas1a* mRNA were measured by qRT-PCR. (**C**) SVEC4-10 cells were pre-treated with anti-IFNAR1 mAb (MAR1- 5A3) or isotype control antibodies overnight at 37°C. Treated cells were then inoculated with CHIKV-AF at a MOI of 1 and collected at 24 hpi for analysis of intracellular E2 expression as described in **Fig 3**. Statistical analysis: Two-way ANOVA test with Šídák’s post-test; \**P* < 0.05 \*\*\*\**P* < 0.0001. Dotted horizontal lines on graphs indicate limit of detection for the assay.

**Figure S7.**
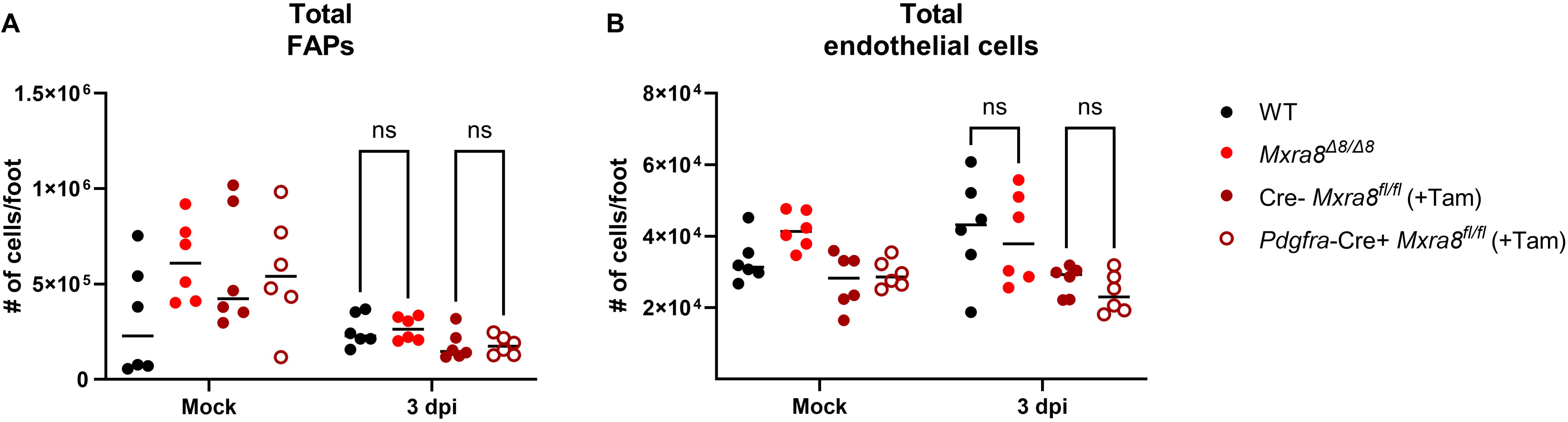
Number of FAPs and endothelial cells isolated from mock and CHIKV-infected mice. Cohorts of 4-week-old C57BL6/J WT, *Mxra8*^Δ*8/*Δ*8*^, tamoxifen (Tam)-treated *Pdgfra-Cre^ERT2^, Mxra8^fl/fl^,* or Tam-treated Cre*-* mice (n = 6 mice per group, 2 experiments) were inoculated with 10^3^ FFU of CHIKV-AF in the left rear footpad or mock-infected. At 3 dpi, mice were administered brefeldin A via intraperitoneal injection 6 h prior to sacrifice. Ipsilateral feet were harvested and digested to release stromal cells. Cell suspensions were incubated with anti-CD45, anti-CD31, anti-CD29, anti-SCA1, anti-VEGFR-2, and anti-VEGF-A antibodies, and the total number of FAPs (**A**) and endothelial cells (**B**) were determined by flow cytometry. Statistical analysis: Two-way ANOVA test with Tukey’s post-test; ns, not significant.

